# Pervasive mislocalization of pathogenic coding variants underlying human disorders

**DOI:** 10.1101/2023.09.05.556368

**Authors:** Jessica Lacoste, Marzieh Haghighi, Shahan Haider, Zhen-Yuan Lin, Dmitri Segal, Chloe Reno, Wesley Wei Qian, Xueting Xiong, Hamdah Shafqat-Abbasi, Pearl. V. Ryder, Rebecca Senft, Beth A. Cimini, Frederick P. Roth, Michael Calderwood, David Hill, Marc Vidal, S. Stephen Yi, Nidhi Sahni, Jian Peng, Anne-Claude Gingras, Shantanu Singh, Anne E. Carpenter, Mikko Taipale

## Abstract

Widespread sequencing has yielded thousands of missense variants predicted or confirmed as disease-causing. This creates a new bottleneck: determining the functional impact of each variant – largely a painstaking, customized process undertaken one or a few genes or variants at a time. Here, we established a high-throughput imaging platform to assay the impact of coding variation on protein localization, evaluating 3,547 missense variants of over 1,000 genes and phenotypes. We discovered that mislocalization is a common consequence of coding variation, affecting about one-sixth of all pathogenic missense variants, all cellular compartments, and recessive and dominant disorders alike. Mislocalization is primarily driven by effects on protein stability and membrane insertion rather than disruptions of trafficking signals or specific interactions. Furthermore, mislocalization patterns help explain pleiotropy and disease severity and provide insights on variants of unknown significance. Our publicly available resource will likely accelerate the understanding of coding variation in human diseases.

## INTRODUCTION

Pathogenic coding variants underlie many human diseases, from rare Mendelian disorders to complex traits, cancer, and neurodegeneration. Rapid advances in genome and exome sequencing technologies have uncovered hundreds of thousands of coding variants associated with these diseases, vastly outpacing our ability to interrogate the effects of coding variation on protein function. This glaring disparity has resulted in two major challenges. On one hand, due to the lack of functional assays, most coding variants in disease genes remain classified as variants of unknown significance (VUS), posing a major roadblock for the clinical interpretation of coding variation^1^. On the other hand, even if a given coding variant is deemed pathogenic, it is often not known how exactly it disrupts protein function^2,3^. Thus, characterizing the functional consequences of coding variants is highly relevant to clinical genetics, understanding disease pathogenesis, and developing novel therapies.

Diseases caused by coding variation are immensely diverse, affecting different cellular processes, tissue types, and developmental stages. Yet, despite this phenotypic diversity, the molecular mechanisms by which coding variants disrupt protein function are much more constrained^2,4^. Mutations often impinge on protein stability, rewire interactions by disrupting existing or introducing new interactions with other biomolecules, or perturb protein subcellular localization. The most common disruptive mechanism is likely the loss of protein stability. Because most proteins are only marginally stable^5^, single amino acid changes can easily destabilize protein folds and promote protein misfolding. Experimental studies have suggested that 30%-60% of all pathogenic missense variants are destabilizing^6,7^. Similarly, pathogenic mutations are enriched in known protein-protein interaction interfaces and can disrupt interactions with nucleic acids or small molecules^7–14^. Our previous work suggests that up to 30% of pathogenic missense mutations disrupt specific protein-protein interactions, i.e. affect some interactions while leaving others unperturbed, indicating that interactome rewiring is a widespread mechanism of pathogenesis^7^.

While large-scale studies and computational approaches have shed light on the frequency of mutations affecting protein stability and interactions, much less is known about mutational effects on protein localization. Correct subcellular localization is fundamental to the function of all proteins, and mislocalization plays a central role in diverse human diseases^2,15–18^. For example, the most common mutation underlying cystic fibrosis, CFTR ΔF508, leads to the retention of the mutant protein in the endoplasmic reticulum (ER) due to ER quality control^19^. Pharmacological correction of CFTR ΔF508 trafficking to the plasma membrane can restore much of CFTR’s activity as a chloride channel, significantly ameliorating symptoms in patients^20^. Similarly, mislocalization of tumor suppressors such as PTEN and BRCA1 and oncogenes such as AKT1 and NPM1 has been causally implicated in tumorigenesis^21–24^. Finally, aberrant localization of proteins to aggregates or phase-separated condensates is a defining hallmark of most neurodegenerative diseases, illustrated by α-synuclein aggregation in Parkinson’s disease and FUS (fused in sarcoma) phase separation in amyotrophic leukodystrophy (ALS)^25^.

Despite many such examples of mislocalized pathogenic variants, computational analyses have suggested that fewer than 2% of pathogenic disease variants are mislocalized^17^. However, this is likely a significant underestimate because accurate prediction of mislocalization is difficult. Protein localization can be regulated by diverse mechanisms, such as compartment-specific sorting signals, protein-protein interactions, cell-state dependent signaling events, ligand binding, or the protein quality control machinery. Disruption of any of these mechanisms could lead to mislocalization. Thus, the full extent to which aberrant protein localization contributes to diverse diseases is still unknown.

Here, we systematically assess how pathogenic coding variants affect protein localization. We use high-content microscopy to characterize the localization pattern of 3,547 missense variants of 1,282 proteins involved in diverse Mendelian disorders and tumorigenesis. Our results reveal that mislocalization is a common phenotype that involves all cellular compartments, with a particularly pronounced role for proteins trafficked through the secretory pathway. Mislocalization equally affects dominant and recessive disorders as well as somatic mutations underlying cancer. We further show that changes in subcellular localization can reveal mechanisms of pleiotropy and help classify variants of unknown significance. We provide the full dataset of images and data as an open-access resource for researchers studying rare diseases and mechanisms of protein trafficking.

## RESULTS

### Systematic analysis of protein localization

To systematically characterize the effect of mutations on protein localization, we used a previously described collection of missense variants (hmORFeome1.1), which contains 3,046 variants across 1,139 genes annotated in the Human Gene Mutation Database (HGMD), in addition to their wild-type, or “reference”, counterparts^7,26^. We complemented this collection with another set of 136 variants (109 missense variants and 27 fusion proteins) in 43 genes encoding protein kinases and chaperone client proteins^27,28^, 295 variants of 115 genes found in cancer genome sequencing projects^29^, and 70 likely non-pathogenic variants of 40 genes identified in exome sequencing studies^7^. Altogether, our collection includes 3,547 variants across 1,282 unique genes, highly enriched for pathogenic and damaging variants based on ClinVar annotations and PolyPhen predictions^30,31^ (**Figures 1A, 1B, S1A**, and **Table S1**). The collection broadly covers variants with autosomal recessive, autosomal dominant, and X-linked inheritance patterns as well as somatic variants and susceptibility alleles (**Figure 1C**).

**Figure 1.**
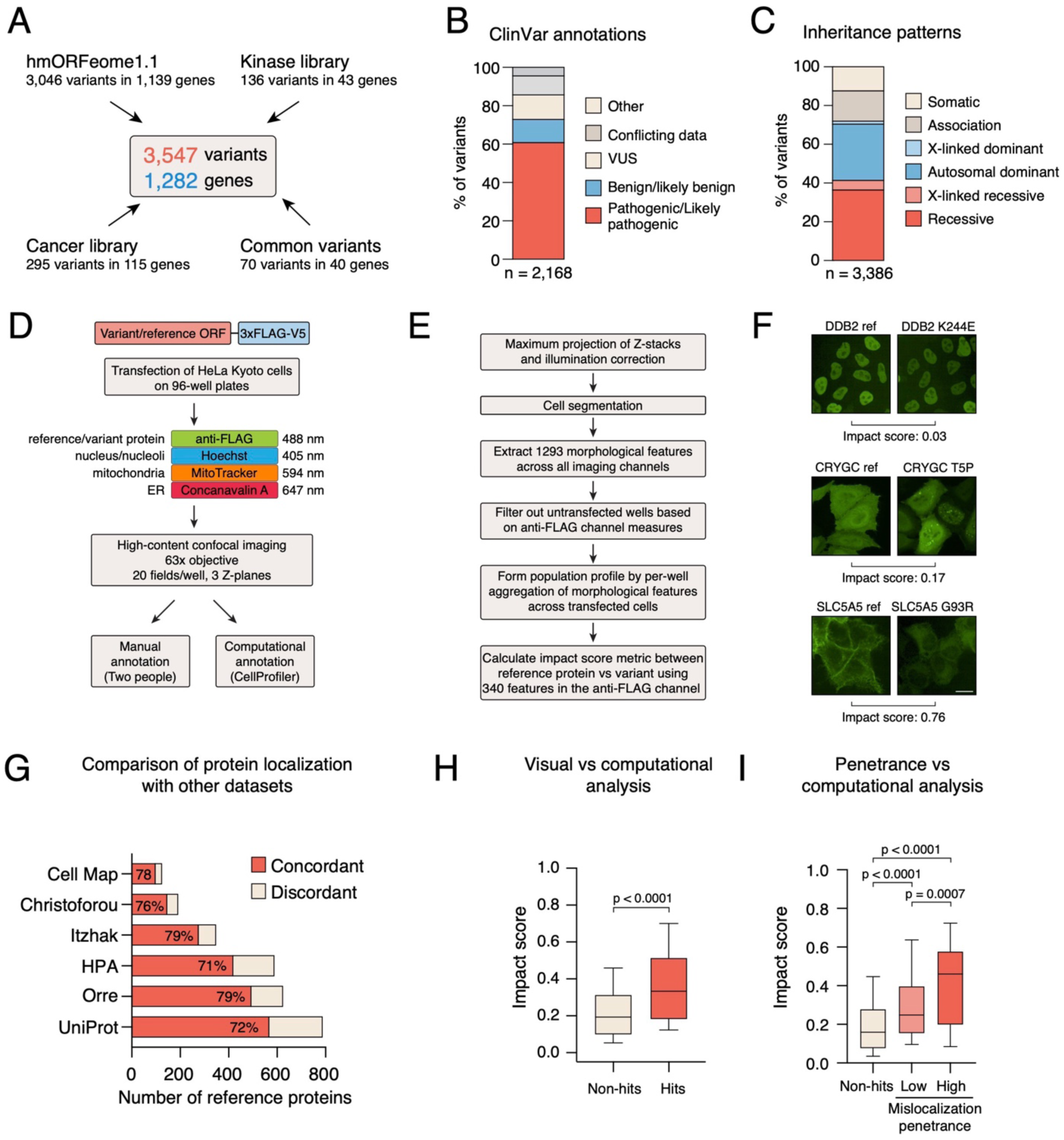
Systematic profiling of subcellular localization of missense variants. **A)** Sources of variants used in this study. **B)** Available ClinVar annotations for the variants used in this study. **C)** Reported inheritance pattern of variants used in this study. **D)** Pipeline for high-content screen for protein localization. **E)** Computational pipeline for analyzing localization patterns and comparison of reference alleles and variants. **F)** Examples of variants with high, medium and low similarity to the reference allele. **G)** Reference protein localization in this study compared to other large-scale studies. **H)** Correlation coefficient of localization patterns between reference alleles and missense variants for visually identified hits and non-hits. Statistical significance was calculated with a Mann-Whitney test. **I)** Correlation coefficient of localization patterns between reference alleles and missense variants for non-hits, low penetrance hits and high penetrance hits. Statistical significance was calculated with ANOVA with Tukey’s correction for multiple testing.

We transfected 3xFLAG-V5 tagged reference and variant constructs into HeLa cells in a 96-well format. After 48 hours, cells were fixed and stained with anti-FLAG antibody and with subcellular markers for nucleus (Hoechst), membranes and endoplasmic reticulum (ER; concanavalin A), and mitochondria (MitoTracker). We used an automated high-content confocal microscope to image 25 fields at three different Z planes of each well using a 63x objective, providing a robust dataset for downstream analysis. All microscopy images were analyzed visually by two independent observers and computationally with a custom CellProfiler pipeline^32^ (**Figures 1D, 1E, and 1F**). Visual and computational annotation both indicated that about 60% of tagged reference and variant proteins were detected in more than 50 cells (**Figure S1B** and **S1C**). Only 12% of proteins were not detected at all (**Figure S1B**), possibly due to low stability or secretion, indicating that our transient transfection protocol provides a robust method for ectopically expressing a large collection of diverse proteins.

We then visually annotated the localization patterns of the detectable reference proteins and their variants, finding the localizations to be generally concordant with publicly available annotations. Excluding secreted proteins that cannot be accurately detected by imaging, over 70% of the detectable reference proteins were localized to at least one of the compartment annotations in six different data sources, including UniProt and the Human Protein Atlas^33^, as well as four independent mass spectrometry-based approaches (**Figure 1G**)^34–37^. These results are in line with previous experiments comparing transfection of epitope-tagged proteins and immunofluorescence with antibodies against endogenous proteins^38^. Notably, only the Itzhak et al. study used HeLa cells, suggesting protein localization is not generally cell-type dependent. To test this more directly, we randomly selected 118 reference and variant proteins and transiently transfected them into HeLa, RPE1, and U2OS cells. Although some proteins showed subtle differences in localization between cell lines, all tested reference or variant proteins were either highly similar or identical when comparing all three cell lines (**Figure S1D** and **S1E**). Variant mislocalization could also be detected in all three cell lines (**Figure S1F**). We conclude that ectopic expression of epitope-tagged proteins is generally a reliable method for detecting protein localization and is not overly influenced by the cell type.

The most common localization compartments for reference proteins were cytoplasm (32%), endoplasmic reticulum (26%), nucleoplasm (14%), mitochondria (8%), plasma membrane (7%), and Golgi apparatus (7%) (**Figure S1G**). However, the majority (54%) of reference proteins localized to multiple compartments. The ER and mitochondria contained the most specifically-localized proteins, while the plasma membrane, Golgi apparatus, vesicles, cytoplasm, and nucleus contained more multi-localizing proteins (**Figure S1H**). These patterns are concordant with previous studies in both human and yeast cells^33,39^.

### Widespread mislocalization of pathogenic missense variants

We then examined the extent to which variants found in humans change the localization of their respective reference proteins. Variants that had a visually different localization pattern than the reference (as assessed by either of the two observers) were classified as high penetrance (more than 50% of cells showed a phenotype) or low penetrance (fewer than 50% of the cells with differing phenotype). We complemented the visual analysis with a computational analysis based on 340 morphological features extracted by a custom CellProfiler pipeline (Figure 1E)^32^. The wild-type/mutant impact was assessed by measuring the distance between the morphological representation of each wild-type/mutant morphological profile (see Method Details). Negative correlation coefficient between well-level preprocessed and aggregated morphological profiles of wild-type/mutant pairs was scaled to a [0,1] range and used as the measure of distance (impact score). All variants that were scored as mislocalized by either visual examination or computational analysis were re-arrayed and tested with a second, independent round of transfection and immunofluorescence. Variants that passed the secondary validation round (which used the same analysis parameters) were considered final mislocalized hits. Highlighting the complementary nature of our approaches, computational analysis detected several subtle changes in localization that were missed by manual observation, whereas manual annotation identified cases where only a limited number of cells expressed the reference or variant protein or cases in which the phenotype was visible in only a fraction of cells. Even so, the approaches were generally in good agreement: hits identified visually had a significantly higher impact score relative to their reference counterpart than non-hits (Figure 1H). Moreover, higher-penetrance hits (as defined manually) had a more dramatic impact on computationally-determined localization than lower-penetrance hits and non-hits (Figure 1I).

Our screen identified 250 (11%) confirmed mislocalized variants out of 2,280 variants that were detected by imaging (Figure 2). These variants represented 152 distinct genes; 16% of genes had at least one mislocalized missense variant. However, nearly 40% of genes for which we assayed four or more variants had at least one mislocalized variant (**Figure S2A**). Thus, missense variation can affect the localization of a substantial portion of the proteome.

**Figure 2.**
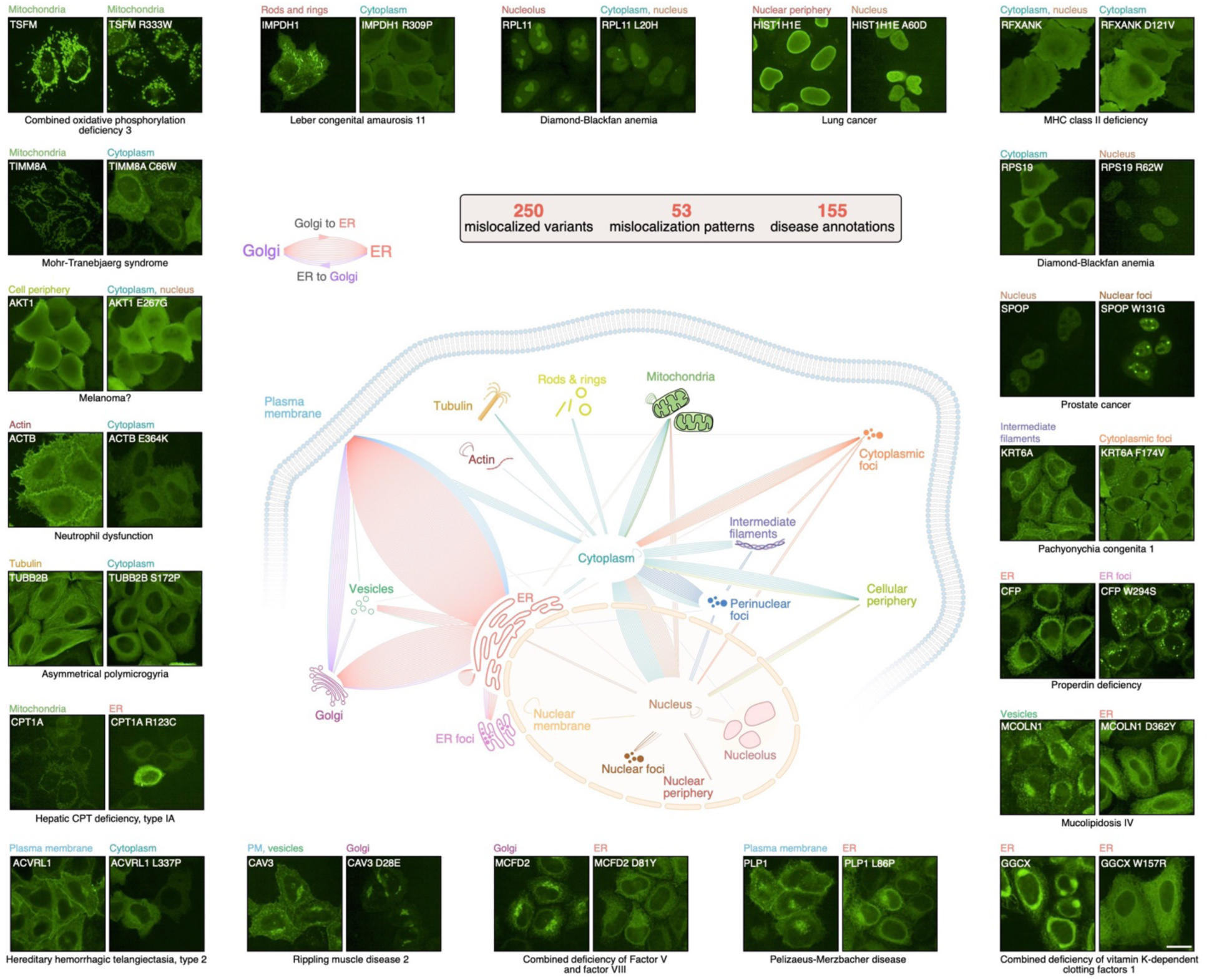
Mislocalization map of missense variants. The overall pattern of mislocalization is shown in the center. Each line represents a mislocalized variant. The color of the line indicates destination compartment (i.e. mislocalization compartment). Examples of mislocalized variants for different categories are shown with manual annotation and the associated disease phenotype. Scale bar, 20 µm.

Mislocalized variants were not equally distributed among genes: if a gene had one mislocalized variant, it was significantly more likely to have multiple such variants than expected by chance (p < 0.0001, Fisher’s exact test; Figure 3A). Thus, mislocalization disproportionately affects some gene products while sparing others. We observed a similar trend with localization patterns. We identified 53 different localization categories from one compartment to another, representing all major cellular compartments (Figure 2). We found that some compartments were significantly more likely to be involved in mislocalization. For example, proteins normally localized to the plasma membrane, Golgi, and vesicles were enriched in mislocalized variants, whereas those localized to the cytoplasm or mitochondria were depleted (Figure 3B). In total, 59% (148/250) of mislocalization events included the secretory pathway (ER, Golgi, plasma membrane, or vesicles) as the reference or variant localization, although these proteins represented only 36% of the library (p < 0.0001; Fisher’s exact test; Figure 3C). These results are consistent with the well-established role of ER in protein quality control^40^. Another class of proteins that displayed frequent mislocalization were cytoskeletal proteins such as keratins, actin, and tubulin, whose assembly into long filaments is central to their function (Figures 3B and **3D**).

**Figure 3.**
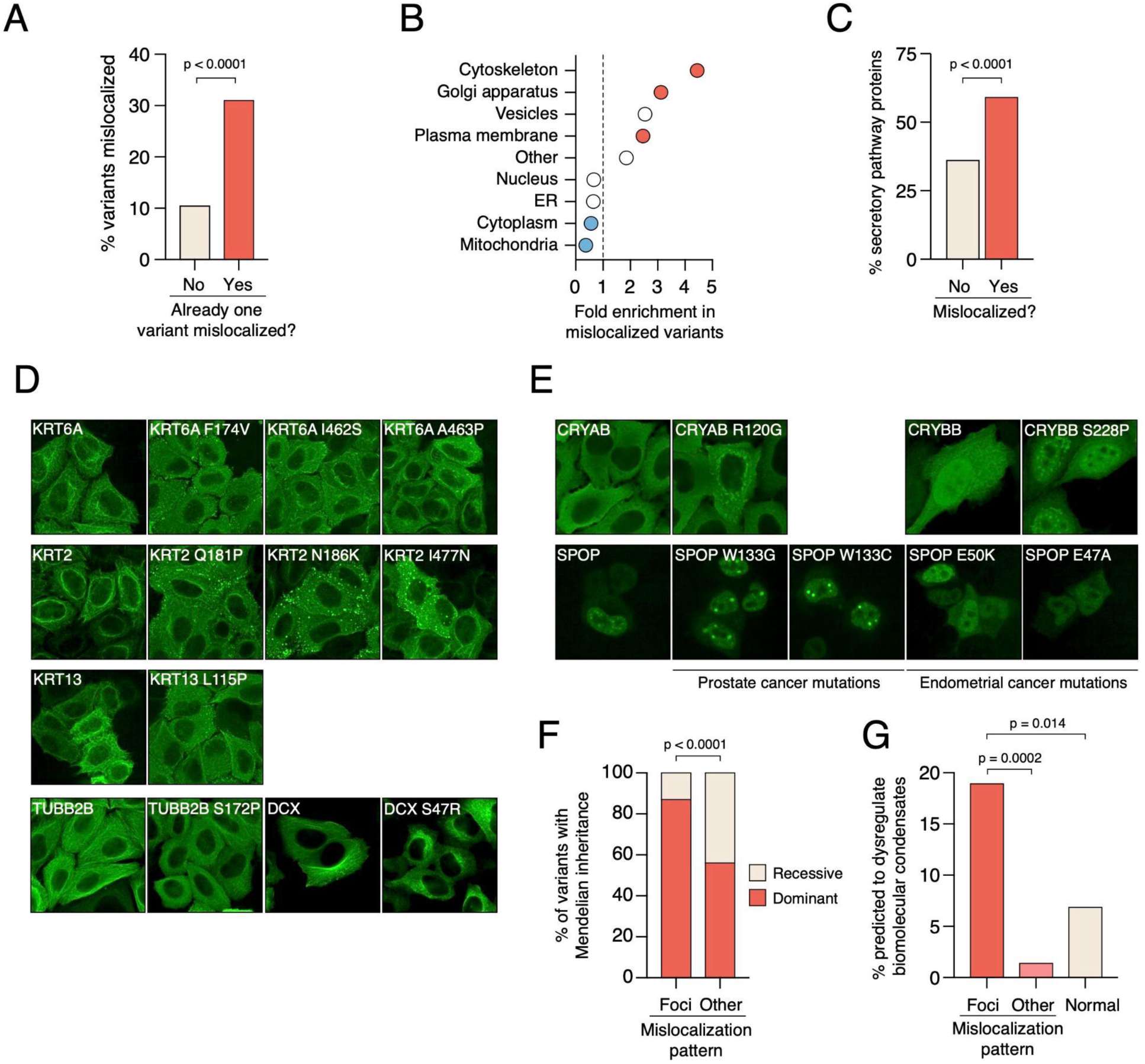
Mislocalization and cellular compartments. A) Mislocalization affects some genes more than others. Fraction of mislocalized variants for all genes compared to those genes for which already one variant is mislocalized. Statistical significance was calculated with Fisher’s exact test. **B)** Mislocalization affects some compartments more than others. Relative enrichment of mislocalized variants by localization of the reference protein. Red and blue circles represent compartments from where significantly more (red) or fewer (blue) variants are mislocalized. **C)** Mislocalized variants are enriched in proteins normally localized to the secretory compartment. Statistical significance was calculated with Fisher’s exact test. **D)** Examples of mislocalized variants of cytoskeletal proteins. Top, keratin proteins forming distinct punctae. Bottom, mislocalized variants of tubulin and doublecortin, a microtubule-associated protein. **E)** Examples of missense variants forming distinct foci. Missense variants of SPOP associated with prostate cancer form more foci than the reference protein, whereas coding variants associated with endometrial cancer do not form foci. **F)** Inheritance pattern of mislocalized variants forming distinct foci and those with other localization patterns. Statistical significance was calculated with Fisher’s exact test. **G)** Comparison of mislocalization results from this study and from Banani et al. for variants predicted to dysregulate biomolecular condensates. Statistical significance was calculated with Fisher’s exact test with Bonferroni correction for multiple hypotheses.

One prominent class of mislocalization was the formation of discrete foci or clusters by 14% (34/250) of the hits, representing 34 distinct proteins (Figures 2 **and 3E**), across a range of baseline reference locations. Among the focus-forming variants were known aggregation-prone proteins, such as germline variants CRYAB R120G and CRYBB1 S228P^41,42^ and multiple somatic variants of the cancer driver SPOP (Figure 3E). Two prostate cancer associated variants of the SPOP E3 ligase (W133G and W133C) formed visibly larger and rounder foci in the nucleus than the reference protein (Figure 3E). These larger foci likely represent membraneless granules previously reported for SPOP^43^. Interestingly, two endometrial cancer-associated SPOP mutants (E47A and E50K) did not form any foci (Figure 3E), indicating that oncogenic variants associated with different tumor types show distinct localization phenotypes. SPOP localization is affected by its substrates^44^, and the endometrial and prostate cancer associated variants have distinct effects on SPOP substrates such as BRD3 and BRD4^45^, likely explaining the difference in localization.

We then asked whether variants whose mislocalization involved loss or gain of foci had features distinguishing them from other mislocalized variants. We found distinct features that were consistent with these variants functioning through a toxic gain-of-function mechanism due to aggregation or condensate dysregulation. First, focus-forming and focus-dissolving variants were enriched for those that function in a dominant manner, compared to other mislocalized variants (Figure 3F). Second, they were significantly enriched in mutations predicted to dysregulate condensate formation, both when compared to other mislocalized variants and to normally localized variants (Figure 3G)^46^. Our results suggest that aggregation and/or condensate dysregulation are common mechanisms of variant mislocalization and human disease pathogenesis, consistent with recent reports^18,46^.

### Features associated with variant mislocalization

To understand the causes underlying protein mislocalization, we compared features of mislocalized variants to normally localized variants. Overall, mislocalization occurred at a similar rate between dominant and recessive variants and between germline and somatic variants (**Figures S2B** and **S2C**), indicating that protein mislocalization broadly affects all types of variation. However, mislocalized variants were significantly enriched in pathogenic variants and depleted of benign variants, as assessed by ClinVar and Polyphen2 (Figures 4A and **4B**), and they had a significantly lower population frequency than normally localized variants (**Figure S2D**)^30,47^. Indeed, not a single common variant from exome sequencing studies (0/36), and only 6% (9/162) of variants annotated in ClinVar as benign or likely benign were mislocalized. In contrast, 16% (135/822) of those annotated as pathogenic or likely pathogenic showed a distinct localization pattern from the reference protein (p < 0.001, Fisher’s exact test; Figure 4C). This pattern was even more pronounced for proteins normally localized to the secretory pathway: 23% of pathogenic or likely pathogenic variants in these proteins were mislocalized (**Figure S2E**).

**Figure 4.**
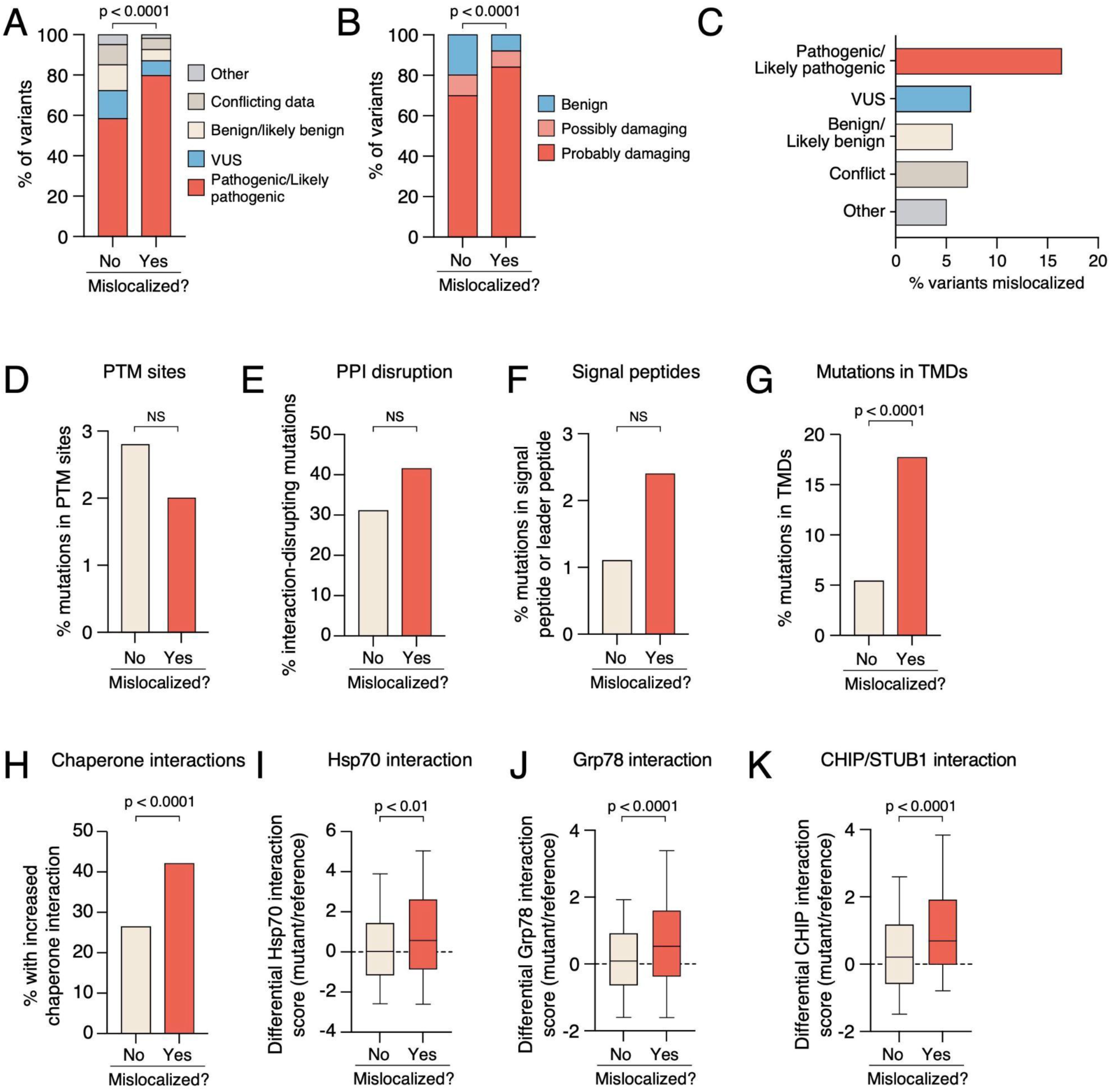
Features associated with mislocalization. A) Mislocalized variants are enriched in pathogenic and likely pathogenic variants. **B)** Mislocalized variants are predicted to be more damaging by Polyphen-2. **C)** Pathogenic and likely pathogenic variants are mislocalized more often than benign or likely benign variants. **D)** Mutations causing mislocalization are not enriched in post-translational modification sites. **E)** Mutations causing mislocalization do not disrupt protein-protein interactions more often than mutations leading to normal localization, as assessed by yeast two-hybrid assay and validated by orthogonal assays^7^. **F)** Mutations causing mislocalization are not enriched in signal peptides. **G)** Mutations causing mislocalization are highly enriched in transmembrane domains. **H)** Mislocalized variants interact more with chaperones and quality-control factors than normally localized variants, as determined by LUMIER assays^7^. **I-K)** Comparison of chaperone and quality control factor interactions of reference proteins and mislocalized or normally localized proteins for Hsp70/HSPA8 **(I)**, Grp78/HSPA5 **(J)**, and Chip/STUB1 **(K).** Statistical significance was calculated with a chi-square test (panels A and B), Fisher’s exact test (panels D-H), and Mann-Whitney test (panels I-K)

We identified nine mislocalized variants classified as benign or likely benign in ClinVar. However, further investigation revealed that at least six of these have been shown to have altered activity or localization in functional assays^7,48–53^, corroborating our results. These variants may be misannotated in ClinVar or cause partially penetrant phenotypes that do not manifest themselves in all individuals. Alternatively, it is possible that these missense mutations functionally affect the protein without having an influence on the organismal phenotype, as has been observed for some protein/protein interaction disrupting variants^54^.

Aberrant protein localization can be caused by mutations that disrupt post-translational modification sites, specific protein-protein interactions, or trafficking signals such as the nuclear localization signal or the signal peptide. Alternatively, it can be caused by mutations that disrupt protein stability, leading to protein misfolding and trapping in intermediary compartments such as the ER ^2^. To understand the relative contributions of these alternatives, we investigated the characteristics of the missense mutations leading to mislocalization.

Only five of the 250 mislocalized variants contained a mutation in a known post-translational modification site (Figure 4D). Notably, two of these variants impact a phosphoserine site in proteins normally localized to microtubules (TUBB2 S172P and DCX S47R) and have been previously reported to affect protein localization and function^55,56^. We observed similar phenotypes for these variants (Figure 3D), suggesting that our approach could capture variant effects on PTM sites. However, overall mislocalized variants were not enriched for mutations in PTM sites, including disulfide bridges (Figures 4D and **S2F**).

Disruption of protein-protein interactions (PPIs) similarly was not a major cause of mislocalization patterns, although again our strategy identified known examples. PPIs can orchestrate the trafficking of proteins to specific compartments^57^. We found that MCDF2 D81Y, which underlies Combined Deficiency of Factor V and Factor VIII (OMIM 613625), was mislocalized from the Golgi apparatus to the ER (Figure 2). Mislocalization of MCFD2 D81Y has been attributed to the loss of interaction with LMAN1, a cargo receptor-like protein that cycles between the ER and Golgi apparatus^58^. To examine the overall contribution of protein-protein interaction patterns on protein localization, we assessed if mislocalized variants were more likely to exhibit PPI perturbations. We therefore compared our imaging results with a previously generated dataset of interactions for the hmORFeome 1.1 collection^7^. There was no significant difference in the frequency of interaction-disrupting mutations between mislocalized and normally localized variants (Figure 4E), suggesting that disruptions in specific PPIs are not a major cause of mislocalization. Consistent with this notion, mutation frequencies in signal peptides and mitochondrial leader peptides, which are recognized by specific PPIs, were also similar between the two groups of variants (Figure 4F). Thus, while the examples highlighted here indicate that our platform could identify known cases of mislocalization due to disruptions in PPIs and PTMs, overall, these do not seem to significantly contribute to protein mislocalization.

In contrast to mutations affecting PTMs, signal peptides, and specific protein-protein interactions, mislocalized variants were highly enriched in mutations that interfere with protein folding or insertion of transmembrane domains into the membrane. Insertion into the lipid bilayer is primarily driven by interactions between aliphatic hydrophobic residues in a transmembrane domain (TMD) and the lipid hydrocarbon chains in the lipid membrane. Nearly 20% of mislocalized variants had a mutation in an annotated TMD, in contrast to only 5% of normally localized variants (p < 0.0001; Fisher’s exact test; Figure 4G). Moreover, TMD mutations in mislocalized variants were significantly enriched in non-conservative substitutions compared to normally localized variants (**Figures S2G** and **S2H**).

To assess the contribution of protein stability, we compared how mislocalized and normally localized variants interacted with chaperones and other cellular quality control factors, which was characterized in our previous study^7^. Mislocalized variants were significantly more likely to interact more with one or more quality control factors than normally localized variants (Figure 4H). However, the pattern was not identical between different quality control factors. For example, mislocalized variants interacted significantly more with Hsp70 family chaperones in the cytoplasm (Hsc70) and ER (Grp78) but not with Hsp90 family chaperones (cytoplasmic Hsp90 and ER-resident Grp94) (Figures 4I-J and **S2I-J**). Consistent with this, Hsp70 cochaperones STUB1 and BAG2 also interacted more with mislocalized variants (Figures 4K and **S2K**). The difference between Hsp70 and Hsp90 interactions reflects the functional differences between these conserved chaperone families. Hsp70 is an early-stage chaperone that promotes *de novo* folding and trafficking of its clients, whereas Hsp90 acts at a later stage and with a more limited set of clients^28,59,60^. In any case, these results establish that protein instability, rather than loss of specific protein/protein interactions, is a major factor driving protein mislocalization.

### Imaging provides insights into mechanisms of variant pathogenicity and disease severity

Disruptions of specific protein/protein interactions can explain pleiotropy, i.e. where two mutations in the same gene cause distinct diseases^7^. We hypothesized that protein localization could underlie pleiotropy in a similar manner. We therefore focused on cases where two variants of a gene had distinct localization patterns and asked whether the associated phenotype annotations were concordant or discordant. Indeed, variant pairs that were differentially localized were significantly enriched for discordant disease annotations compared to pairs that were similarly localized (Figure 5A).

**Figure 5.**
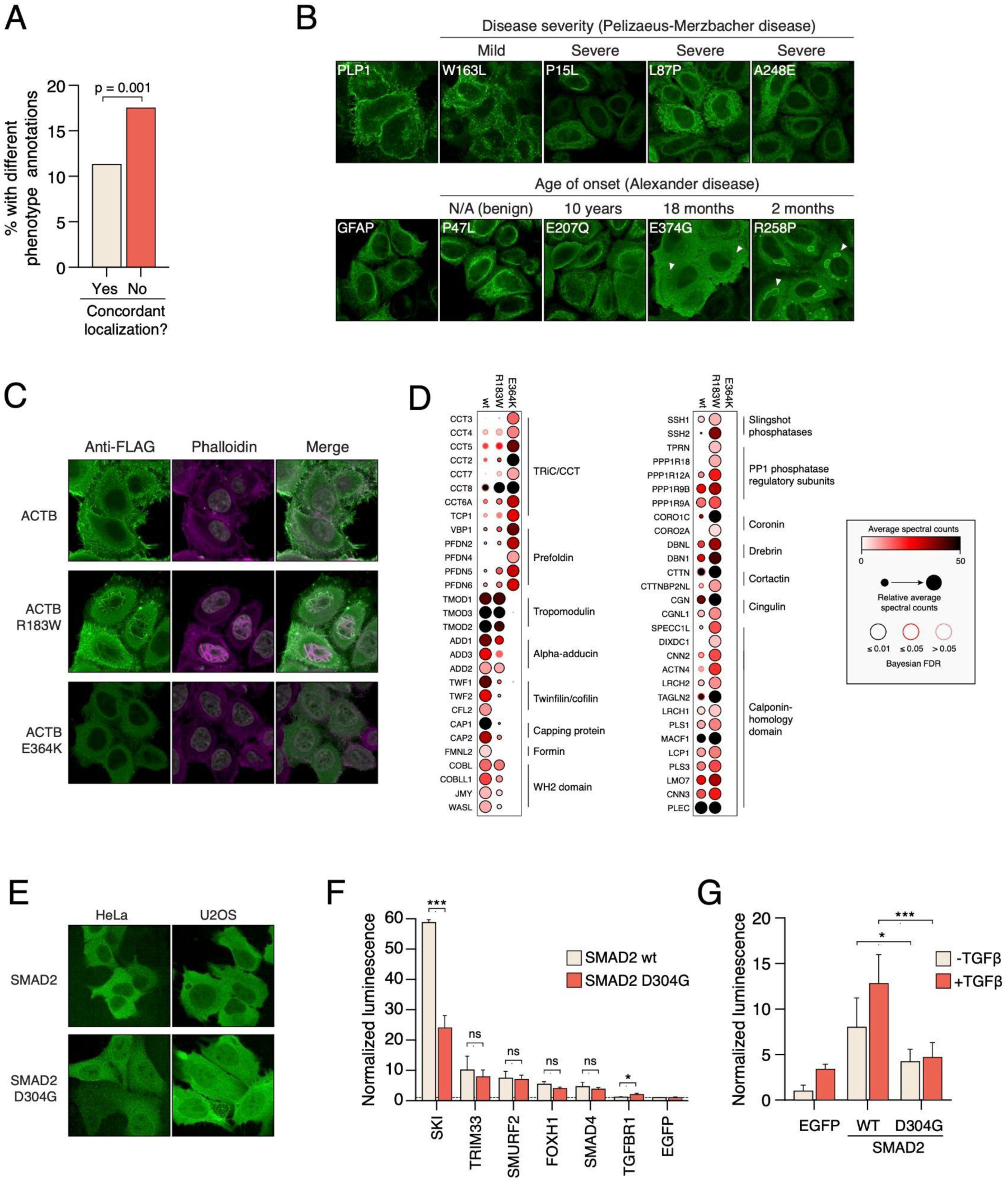
Mislocalization and disease phenotypes. A) Variants of the same gene that have a distinct localization pattern are more often associated with distinct disease phenotypes than variants that are similarly localized. Statistical significance was calculated with Fisher’s exact test. **B)** Top, loss of membrane localization of PLP1 variants is concordant with disease manifestation. Bottom, loss of intermediate filament staining and appearance of distinct punctae (arrowheads) with GFAP variants correlates with age of onset. **C)** Distinct localization of beta-actin variants underlying different diseases. Filamentous actin staining with phalloidin (magenta) shows distinct patterns with wild-type actin and each mutant. **D)** Proximity interactomes of wild-type actin and R183W and E364K variants were determined by BioID in HEK293 cells. The graph shows selected interactions; the full dataset is available in Table S1 **E)** Mislocalization of SMAD2 D304G variant from the nucleus to the cytoplasm. Wild-type and mutant SMAD2-3xFLAG-V5 constructs were transfected into HeLa and U2OS cells and stained with anti-FLAG antibody (green). **F)** SMAD2 D304G interacts less with the transcriptional co-regulator SKI and more with the TGFβ receptor TGFBR1. Indicated 3xFLAG-V5 tagged constructs were co-transfected into HEK293T cells with Nanoluc-tagged wild-type SMAD2 or the D304G variant, and interaction was assayed with LUMIER assay. Statistical significance was calculated with ANOVA with Tukey’s correction for multiple hypotheses. ***, p < 0.001, *, p < 0.05. **G)** SMAD2 D304G is a weaker transactivator than wild-type SMAD2. Indicated constructs were co-transfected with 3TP-lux reporter and Nanoluc control into MDA-231 cells and the cells were treated with vehicle control or TGFβ. Transactivation activity measured with luciferase assay. The ratio between Firefly and Nanoluc luminescence was normalized to EGFP control with vehicle treatment. Statistical significance was calculated with ANOVA with Tukey’s correction for multiple hypotheses. ***, p < 0.001, *, p < 0.05.

To investigate if localization patterns could similarly provide insights into disease severity, we surveyed the literature for mislocalized variants that had annotations for disease severity or age of onset. We found such annotations for PLP1 and GFAP variants, underlying distinct leukodystrophies. PLP1 (myelin proteolipid protein) mutations cause Pelizaeus-Merzbacher disease, a demyelinating disorder that is manifested as a spectrum of symptoms contingent on the genotype^61^. Gain-of-function mutations in GFAP (Glial fibrillary acidic protein) cause autosomal dominant Alexander disease, which has a highly variable disease presentation and age of onset^62^. In both cases, we observed a strong correlation between disease severity and the extent of mislocalization. PLP1 W163L, which is associated with a very mild form of Pelizaeus-Merzbacher disease^63^, showed slightly decreased plasma membrane staining compared to the reference protein, whereas three variants underlying severe (connatal) disease lost all plasma membrane localization (Figure 5B). This concordance between subcellular localization and patient phenotype is consistent with a previous study with two other PLP1 variants^64^. GFAP variant localization ranged from reference-like intermediate filament localization to diffuse cytoplasmic staining to prominent cytoplasmic punctae, likely reflecting pathogenic aggregation^62^ (Figure 5B). The localization pattern correlated with the age of onset of the disease: the E207Q variant associated with the oldest age of onset showed only a partial diffuse localization, whereas the R258P variant with the youngest age of onset was localized to distinct punctae. The third pathogenic variant (E374G), associated with an intermediate age of onset, localized mostly to the cytoplasm, with some cells showing weak punctate staining. Thus, our results suggest that protein mislocalization is associated with both pleiotropy and disease severity.

We then searched for examples where our imaging results could provide insight into the potential pathogenic mechanisms of variants with distinct localization patterns. Among our hits were two β-actin (ACTB) variants, R183W and E364K. In humans, these variants lead to distinct phenotypes: R183W was found in a patient with developmental malformations, deafness, and delayed-onset dystonia^65^, whereas E364K is associated with neutrophil dysfunction^66^. *In vitro* assays with purified proteins have not revealed dramatic changes in these variants’ thermal stability compared to the wild-type protein. However, the two mutations have similar but modest effects on the polymerization and ATP hydrolysis activity of actin^67^. Moreover, *in vitro* studies with the E364K mutant have provided conflicting evidence as to its effects on folding and profilin binding^66–68^, and this variant is classified as a VUS in ClinVar.

In contrast to these *in vitro* assays, imaging revealed striking differences in the localization of ACTB R183W and E364K compared to wild-type β-actin. ACTB R183W formed remarkable filaments overlapping the nucleus, whereas ACTB E364K showed reduced protein levels and did not form any filaments (Figure 5C). Staining filamentous actin with phalloidin corroborated these results: cells expressing the R183W variant displayed prominent actin filaments, whereas E364K-expressing cells had less prominent F-actin staining than those expressing wild-type actin.

To gain more insight into how these mutations lead to distinct phenotypes, we characterized the interactomes of wild-type ACTB and the two variants with proximity-dependent biotinylation (BioID). We generated stable tetracycline-inducible HEK293 cell lines expressing each construct fused to the FLAG epitope and the abortive biotin ligase BirA*, which promotes biotinylation of proximal proteins. We first analyzed construct localization by anti-FLAG immunofluorescence and streptavidin staining after biotin treatment. Similar to HeLa cells, ACTB R183W and E364K showed prominent differences from each other and from the wild-type β-actin (**Figure S3A**). E364K was again expressed at lower levels than wild-type β-actin and localized in the cytoplasm in a diffuse manner. In contrast, the R183W variant was localized to membrane-proximal regions like wild-type β-actin, but it also formed large cytoplasmic foci (**Figure S3A**). The difference in ACTB R183W localization between HEK293 and HeLa cells is likely due to cell type differences in actin dynamics and regulation in these cell lines. However, the prominent differences in R183W and E364K localization in both systems strongly suggest distinct functional consequences of these mutations.

Consistent with our imaging results, the two variants showed clear differences in their proximity interactions (Figure 5D and **Table S1**). The ACTB E364K interaction pattern was consistent with a loss-of-function phenotype. This mutant lost proximal interactions with virtually all proteins interacting with wild-type β-actin, but associated more with subunits of the TRiC/CCT chaperonin and prefoldin, which are key chaperones regulating actin folding^69^. This is also consistent with our previous finding that ACTB E364K interacts more than wild-type β-actin with several cellular chaperones^7^. Thus, E364K very likely represents a loss-of-function β-actin variant due to deficient folding, which is not readily seen with *in vitro* assays. Our results also suggest that E364K should be re-classified as a true pathogenic variant.

ACTB R183W, in contrast, showed a complex pattern of changed proximal interactions. While it interacted with some known actin regulators (e.g. tropomodulins) to a similar extent as wild-type β-actin, most proximal interactions were either increased or decreased. For example, the variant associated less with proteins that bind the barbed end of the actin filament, such as capping proteins (CAP1 and CAP2), formins (FMNL2 and FMNL3), and WH2 domain proteins (JMY, WASL). On the other hand, it associated more strongly with proteins that associate with the side of filamentous actin and/or promote actin bundling and cross-linking, including alpha-actinin and other proteins with calponin-homology domains, regulatory subunits of protein phosphatase PP1, and tropomyosin^70^. This interaction pattern suggests that the R183W mutation affects actin dynamics by promoting the association of filament-stabilizing proteins and disrupting interactions with factors promoting polymerization or depolymerization. The prominent differences between E364K and R183W variants in localization and proximity interaction partners likely explain the distinct disease manifestations associated with these mutations.

Next, we investigated if imaging could provide functional insights into variants identified in cancer genome sequencing studies. Our imaging screen identified a subtle but reproducible phenotype in a mutant variant of the SMAD2 transcription factor, SMAD2 D304G. The mutant variant localized more to the nucleus than wild-type SMAD2, which was mostly cytoplasmic (Figure 5E). We observed a similarly subtle but reproducible phenotype in U2OS cells, corroborating the original screen results (Figure 5E).

SMAD2 is a key transcription factor in the TGFβ pathway. Loss-of-function mutations or deletions in SMAD transcription factors and TGF-β receptor genes occur in a significant fraction of esophageal, gastric, colorectal, and pancreatic adenocarcinomas^71^. Upon TGFβ pathway stimulation, SMAD2 interacts with SMAD4, translocates to the nucleus, and activates TGFβ responsive genes. The D304G mutation is located in the C-terminal MH2 domain, which regulates interactions of SMAD2 with other SMADs, transcriptional cofactors and other cellular factors^71^. We therefore investigated if the mutation might disrupt some of these interactions. To this end, we used the LUMIER assay, which has previously been used to study SMAD2 interactions^72^. We co-transfected Nanoluc-tagged reference SMAD2 or D304G with 3xFLAG-tagged SMAD2 interaction partners and measured luminescence after anti-FLAG pulldown. Wild-type and mutant SMAD2 interacted equally well with the known SMAD2 interactors TRIM33, SMURF2, FOXH1, and SMAD4 (Figure 5F). However, the D304G mutant had a severely reduced interaction with the transcriptional regulator SKI and a slightly increased interaction with the TGFβ receptor TGFBR1. Thus, the mutation selectively affects some SMAD2 interactions while having no effect on others.

To assess the effects of the D304G mutation on SMAD2 activity, we assayed its ability to activate a TGFβ reporter gene^73^. We co-transfected 3xFLAG-tagged wild-type SMAD2 or D304G with the 3TP-lux reporter and Nanoluc control plasmid into MDA-231 cells and measured reporter activity with or without TGFβ stimulation. Wild-type SMAD2 robustly activated the reporter in control conditions, and this was boosted by TGFβ treatment (Figure 5G). In contrast, SMAD2 D304G activated the reporter much less and it did not respond to TGFβ (Figure 5G), indicating that the mutation disrupts SMAD2’s transactivation potential and TGFβ responsiveness. The loss of transactivation capability suggests that the D304G variant is phenotypically a loss-of-function mutation, thereby potentially contributing to tumorigenesis. More generally, these results show that protein localization can be used to prioritize variants of unknown significance for functional studies.

## DISCUSSION

This study represents the first large-scale, publicly available map of the impact of human coding variants on protein localization. Aside from serving as a freely available resource for researchers interested in each variant, reference gene, or associated human disorder, our work answers fundamental questions about the frequency, characteristics, and mechanisms of mislocalization in human disease..

Our work firmly establishes that protein mislocalization is a common result of pathogenic missense variation in diverse disease genes. At least one in six pathogenic or likely pathogenic variants are mislocalized, rising to at least one in four for variants in proteins trafficked through the secretory pathway. Thus, mislocalization is nearly as common a molecular phenotype of pathogenic missense variants as loss of protein stability or loss of protein-protein interactions. Moreover, mislocalization is equally involved in variants underlying recessive and dominant diseases, as well as germline and somatic variants, illustrating the central role of aberrant protein trafficking and localization in disease pathogenesis.

### Compartments of mislocalization

While variant mislocalization affected all subcellular compartments, it was particularly common in the secretory pathway. This is consistent with the compartmentalized nature of protein quality control in the secretory pathway. Mutants that disrupt the folding of proteins trafficked through the secretory pathway are often retained in the ER or Golgi before being targeted for degradation via the proteasome, autophagy, or ER-phagy^40^. Thus, any mutations that interfere with protein folding in the ER or insertion into the membrane will likely lead to mislocalization. Studies with model substrates in yeast and human cells have delineated multiple pathways that regulate protein quality control in the secretory pathway. However, although the pathways are well known, it is still poorly understood how individual substrates are recognized by one or another pathway^40^. We believe that the large-scale collection of mislocalized secretory pathway variants will be a valuable resource to more comprehensively characterize these pathways in a disease-relevant setting.

We also observed that 1.5% of all tested variants (corresponding to 14% of all mislocalized variants) formed distinct punctate structures compared to the reference protein. These included known aggregating variants in lens crystallins and cytoskeletal keratins as well as variants known to regulate protein partitioning to biomolecular condensates. Importantly, variants affecting punctate structures were significantly enriched for those that are predicted to modulate condensation properties of proteins and for autosomal dominant inheritance pattern. These independent lines of evidence suggest that many of the newly discovered variants could be pathogenic due to their propensity to interfere with biomolecular condensates in a gain-of-function or dominant negative manner. We suggest that such focus-forming variants in our study be prioritized for further studies to characterize their mechanism of action in molecular detail. Of note, although the absolute frequency of focus-forming variants was relatively low, the highly charged 3xFLAG-V5 tag used in our study may interfere with condensate formation. It is known that different tags can have distinct effects on the phase separation behavior of tagged proteins^74^. Assessing localization patterns with alternative tags such as fluorescent proteins could reveal more variants with altered condensate behavior.

### Causes of mislocalization

Our study suggests that, overall, protein mislocalization is caused more often by mutations disrupting protein stability and folding rather than mutations in specific motifs that regulate protein trafficking or those that interfere with specific protein-protein interactions. This notion was supported by the tendency of mislocalized variants to interact more with cellular quality control factors than normally localized variants, and by the enrichment of mutations in transmembrane domains in mislocalized variants. While there are many exceptions, the trend is consistent with the available target space for mutations. Most proteins have many more residues that contribute to protein stability than those that regulate specific protein-protein interactions involved in trafficking. Our findings also highlight the interconnectivity of molecular phenotypes of disease variants. Missense variants often impinge on multiple facets of protein homeostasis. For example, loss of stability can cause mislocalization, which in turn can affect protein-protein interactions by limiting access to interacting partners. On the other hand, loss of specific interactions can lead to protein instability. Characterizing pathogenic variants with multiple complementary assays can help understand how these processes are connected and provide a means to uncover the root cause of pathogenesis at the molecular level.

### Complementary strategies for variant phenotyping

Our approach to study a few variants but across more than a thousand different genes is highly complementary to deep mutational scanning (DMS) studies. DMS has emerged as a powerful method to systematically characterize the effects of all missense variants on a single protein at a time^1^. However, many DMS studies have employed readouts that are specific to each protein’s function, limiting assay transferability and scalability. On the other hand, “wide mutational scanning” (WMS) provides a phenotypic survey of variants across a wide swath of proteins. Although not all genes or variants are amenable to WMS based on protein localization, combining multiple scalable assays for common phenotypes of pathogenic variants can significantly increase the coverage^75^. Indeed, if our localization results are combined with our previous study for variant stability and protein-protein interactions with the same variant collection^7^, 66% of the genes had at least one pathogenic or likely pathogenic variant with a phenotype in at least one assay. Thus, a relatively limited set of scalable assays for common molecular phenotypes can cover a large fraction of all genes.

Imaging presents several avenues to further increase the sensitivity and throughput of variant phenotyping. We have shown that cell morphological profiling (“Cell Painting”) can predict the impact of coding variants on protein function and distinguish between gain-of-function, change-of-function, and loss-of-function variants of diverse genes^76,77^. Integrating Cell Painting with variant localization in the future could provide an exceptionally sensitive and information-rich platform for variant phenotyping, although more replicates per sample would likely be needed to capture the often-subtle morphological effects of Cell Painting, compared to effects on protein localization. At the same time, moving from arrayed libraries to pooled optical screens enabled by barcoding and *in situ* sequencing^78,79^ could dramatically increase the throughput of variant profiling by imaging. Further improvements in large-scale mutagenesis in human cells and decreases in the cost of DNA synthesis could make it realistic to phenotypically profile all ∼200,000 pathogenic missense coding variants reported in ClinVar, providing an unprecedented resource of variant phenotypes across thousands of rare diseases.

### Mislocalization as a phenotype for drug discovery

The mislocalization phenotypes discovered in our study could serve as starting points for chemical screens for correctors of trafficking defects. Notably, correctors and potentiators of mutant CFTR trafficking, which are now in clinical use, were originally identified in phenotypic screens in cell culture models with ectopically expressed constructs^80,81^. It is highly likely that many other pathogenic variants could be similarly corrected with small molecules. These could directly bind mutant proteins akin to the CFTR correctors or target key nodes regulating protein trafficking. The latter possibility is particularly attractive, as mislocalized variants are significantly enriched in membrane proteins and secreted proteins. This raises the tantalizing possibility that similar therapeutics might be identified that could mitigate an entire class of mislocalizations and, therefore, potentially an entire class of disorders. For example, pharmacological manipulation of the unfolded protein response can promote trafficking of some loss-of-function variants and prevent protein aggregation in the secretory pathway^82–85^. However, the vast majority of mislocalized disease variants remain to be tested for pharmacological rescue. Our comprehensive resource of pathogenic mislocalized variants could act as a scalable platform for the characterization of pharmacological chaperones across hundreds of phenotypes and diseases, providing a springboard for the discovery of novel therapeutics for rare disorders.

## ACKNOWLEDGEMENTS

We would like to thank other members of the Taipale and Carpenter–Singh labs for their help during the project and for comments on the manuscript. This work was supported by a Canadian Institutes of Health Research Project Grant (PJT-153273), Ontario Ministry of Research and Innovation Early Researcher Award (ER15-11-043), and the University of Toronto Connaught Fund award to M.T., National Institutes of Health grants to N.S. (R35 GM137836), S.Y. (R35 GM133658), M.V. (UM1 HG011989), A.E.C. (R35 GM122547), and Susan G. Komen Foundation grant to N.S. (CCR19609287). N.S. is a CPRIT Scholar in Cancer Research with funding from the Cancer Prevention and Research Institute of Texas (CPRIT) New Investigator Grant RR160021. We also thank Spring Discovery for hosting all microscopy images and providing the web interface for analysis.

## AUTHOR CONTRIBUTIONS

Conceptualization: JL, MH, AEC, MT

Investigation: JL, MH, SH, ZYL, DS, CR, WWQ, XX, HAS, PVR, RS, BAC, ACG, SS, AEC, MT

Resources: ZYL, MC, DH, SSY, NS, MT

Formal Analysis: JL, MH, DS, WWQ, HQA, PVR, RS, BAC, SS, AEC, MT

Visualization: JL, MH, MT

Writing – Original Draft: JL, MT

Writing – Review & Editing: JL, MH, DS, ACG, AEC, MT

Supervision: FPR, MC, DH, MV, JP, ACG, SS, AEC, MT

Funding Acquisition: FPR, MC, DH, MV, JP, ACG, SS, AEC, MT

## STAR METHODS

### RESOURCE AVAILABILITY

#### Lead contact

Further information and requests for resources and reagents should be directed to and will be fulfilled by the lead contact, Mikko Taipale.

#### Materials availability

All plasmids and cell lines generated in this study are available from the authors upon request.

#### Data and code availability

All imaging data are available at https://github.com/carpenter-singh-

### lab/2023_LacosteHaghighi_submitted and at

https://app.springscience.com/workspace/utoronto.

## EXPERIMENTAL MODEL AND SUBJECT DETAILS

### Cell culture

Hela Kyoto, U2OS, RPE1, HEK293T, and HEK293 Flp-In TREx cells were cultured in DMEM with 10% FBS and 1% penicillin/streptomycin. MDA-231 cells were cultured in RPMI 1640 medium (Gibco) supplemented with 5% FBS and 1% penicillin/streptomycin. Cells were dissociated with trypsin and all cells were maintained at 37 °C and 5% CO2. Cells were regularly monitored for mycoplasma infection. HeLa Kyoto cells, HEK293T, and HEK293 Flp-In TREx cells were authenticated with STR profiling (GenePrint 24 System, Promega) at The Centre for Applied Genomics, The Hospital for Sick Children, Toronto.

HeLa Kyoto, U2OS, and RPE1 cells used in this study were a gift from the Pelletier laboratory (Mount Sinai, Toronto, Canada). MDA-231 cells were a gift from the Attisano laboratory.

## METHOD DETAILS

### Plasmids

We used the previously described hmORFome 1.1 and common variant collection^7^, a collection of kinase mutant variants^27^, and the Target Accelerator Pan-Cancer Mutant Collection (Addgene Kit #1000000103)^29^.

Entry clones were transferred using Gateway technology into a mammalian expression pcDNA3.1 plasmid containing an N-terminal 3xFLAG-V5 tag (Addgene 87064) for the Target Accelerator Pan-Cancer Mutant Collection, or a pcDNA3.1-based plasmid containing a C-terminal 3xFLAG-V5 tag (Addgene 87063) for all other variants. Inserts were verified by restriction digestion and clones that did not produce the expected digestion pattern were omitted from further analysis.

### Transfection and Immunofluorescence

Cells were seeded on CellCarrier-96 well black, optically clear bottom plates (Perkin Elmer) at a density of 5,000 cells/well and incubated overnight to attach. Plasmids were transfected into HeLa Kyoto cells with X-tremeGENE 9™ (MilliporeSigma) following the manufacturer’s protocol. Two days post-transfection, cells were fixed with 4% paraformaldehyde in culture medium for 20 min at room temperature, followed by three washes in 1xPBS. Cells were permeabilized with 0.1% Triton X-100/1xPBS for 10 min and blocked with 1% BSA/0.1%Triton X-100/1xPBS for 45 min. Cells were incubated with anti-FLAG M2 antibody (1:500, Sigma-Aldrich) diluted in blocking buffer for 1 h at room temperature. Subsequently, cells were washed in 1 x PBS then incubated with Alexa Fluor 488 goat anti-mouse (1:500, Thermo Fisher Scientific), Hoechst 33342 (1:5000, Thermo Fisher Scientific), Concanavalin A, Alexa Fluor™ 647 Conjugate (1:250, Thermo Fisher Scientific) and MitoTracker™ Red CM-H2Xros (100 nM, Thermo Fisher Scientific) diluted in blocking buffer for 1 hour. For phalloidin staining of actin filaments, cells were incubated in Alexa Fluor™ 568 Phalloidin (1:400, Thermo Fisher Scientific) diluted in blocking buffer for 1 hour.

### Image acquisition

For quantitative imaging of stained cells, images were acquired using the Opera Phenix™ screening system (PerkinElmer) using a 63x/1.15 NA water immersion objective in confocal mode. In every experiment twenty-five fields were acquired per well, capturing four fluorescence channels each with Harmony™ high-content imaging software (PerkinElmer).

### Image analysis

Using CellProfiler^31^ software, images were corrected for illumination variation across the field of view, individual cells were segmented, and 1,313 morphological features were measured for each cell across four imaging channels. Then untransfected cells were filtered out based on the mean intensity of the 488 fluorescence channel (FLAG tag). Replicate-level (equivalent to well-level) profiles were formed by aggregation (population-average) of all transfected imaged single cells in each sample well. Features with near zero variance were removed and replicate-level profiles were standardized per plate to have zero mean and unit variance. The similarity between two profiles is measured by the Pearson correlation coefficient (CC) and is used to assess technical replicate reproducibility of the perturbations with more than one technical replicate.

The Impact Score (IS) for each wild-type/mutant pair is then defined as the (1-CC)/2 in which CC is the correlation coefficient between the wild-type and the mutant replicate-level profiles on each same plate. Impact scores were calculated for each wild-type/mutant pair at three levels of feature categories: features relating to the tagged protein channel (n=340), all other channels (n=953), and all four channels (n=1293) combined. An impact score of 0 for a wild-type/mutant pair indicates perfect similarity between the pair, whereas an impact score of 1 indicates that the profiles show opposite patterns. The full data processing and analysis pipeline and workflow is publicly available at: https://github.com/carpenter-singh-lab/2023_LacosteHaghighi_submitted

### Feature analysis

Features of variants were extracted from HGMD, PolyPhen-2, AlphaFold Database, and ClinVar ^26,30,31,86^. Post-translational modifications (PTMs) were extracted from ActiveDriver, PhosphoSite Plus, and UniProt^87,88^.

### Stable cell line production for mass spectrometry

Entry clones were from the collections previously described in Plasmids. Entry clones were transferred using Gateway technology to Flp-In T-REx compatible vector pDEST-pcDNA5-BirA*-FLAG, to express the BirA* at the C-terminus of the bait^89^. All inserts were verified using Sanger sequencing. The resulting constructs were integrated into HEK293 Flp-In T-REx cells using the Flp-In technology (Thermo Fisher Scientific). HEK293 Flp-In T-REx were seeded in 6-well plates and co-transfected the following day with 200 ng bait-BirA*-FLAG construct and 2 μg pOG44 (Invitrogen) with Lipofectamine 2000, as per the manufacturer’s (Invitrogen) protocol. 24 hours later, cells were expanded to a 10-cm dish. Polyclonal cell populations were then selected for 12-15 days with 200 μg/ml hygromycin B. Expression of constructs and biotinylation activity was validated by Western blotting.

### Sample preparation for BioID

Sample preparation and processing was performed in two biological replicates. Cells were grown in 150 mm dishes to roughly 70% confluence, and then gene expression was induced with 1 µg/ml tetracycline. 12 hours later, 50 mM biotin was added to each plate and incubated for another 12 hours. Cells were washed once using 1 x PBS, scraped, pelleted, flash-frozen and kept at 80°C until processing.

Sample processing began with resuspension of cell pellets in ice-cold modified RIPA buffer (50 mM Tris-HCl pH 7.5, 150 mM NaCl, 0.5 mM EDTA, 1 mM EGTA, 1 mM MgCl2, 1% NP40, 0.1% SDS, 0.5% sodium deoxycholate) containing protease inhibitors (1 mM PMSF, 1:500 protease inhibitor cocktail from Sigma-Aldrich, P8340) at a 1:8 pellet weight (g) to lysis buffer volume (ml) ratio. Lysates were sonicated (3 x 5-second bursts with 3 seconds rest in between at 33% amplitude). 1 μL of RNase A (Sigma-Aldrich, R6148) and 1 μL TurboNuclease (BioVision, 9207) were added to each sample, followed by incubation on a nutator/rocker for 20 minutes at 4°C. To further solubilize membranes, appropriate volumes of 10% SDS were added into each sample to bring the SDS concentration up to 0.25%. After 5 minutes of mixing at 4°C, samples were centrifuged at 14,000 rpm for 20 minutes at 4°C. After centrifugation, equal amounts of supernatant from each sample were mixed with 30 μl of pre-washed streptavidin-sepharose beads (GE Healthcare, 17-5113-01) and incubated for 3 hours at 4°C on a nutator. Following affinity purification of biotinylated proteins, the supernatant was discarded, and beads were washed once with modified RIPA buffer containing inhibitors, once with 2% SDS wash buffer (25 mM Tris-HCl pH 7.5, 2% SDS), once with RIPA wash buffer (50 mM Tris-HCl pH 7.5, 150 mM NaCl, 1 mM EDTA, 1% NP40, 0.1% SDS, 0.5% sodium deoxycholate), and finally once with TENN-wash buffer (50 mM Tris-HCl pH 7.5, 150 mM NaCl, 1 mM EDTA, 0.1% NP40). Following those washes, the beads were washed three times with ABC buffer (50 mM ammonium bicarbonate pH 8 in mass spectrometry grade H2O). Streptavidin-sepharose beads were then resuspended in 50 μl of ABC buffer, and 1 μg trypsin (Sigma-Aldrich, T6567) was added to the samples. The samples were incubated overnight at 37°C, followed by addition of 0.5 μg of trypsin for 4 hours at 37°C. After trypsin digestion, the beads were pelleted, and the supernatant was recovered into a fresh 1.5 ml microfuge tube. The beads were rinsed twice in 80 μl of HPLC-grade H2O, and the rinses were pooled with the original supernatant. The pooled supernatants were centrifuged at 16,100 x g for 2 minutes, and all but the bottom 15-20 µl was transferred to a fresh 1.5 mL microfuge tube. The samples were dried with a centrifugal evaporator and stored at −80°C until further processing.

### Mass spectrometry data acquisition

Fused silica (0.75 μm ID, 350 μm OD) capillary columns were pulled with a laser puller and packed in-house with 10–12 cm C18 (Reprosil-Pur 120 C18-AQ, 3 μm, Dr. Maisch HPLC GmbH) in methanol. Columns were equilibrated in buffer A (0.1% formic acid in 2% acetonitrile) before sample loading.

A quarter of each BioID sample was analyzed using the TripleTOF 5600 (AB Sciex) in Data-Dependent Acquisition (DDA) mode. Briefly, 5 μL of digested peptides were loaded at 400 nL/min onto a previously equilibrated HPLC column. The peptides were eluted from the column over a 90-minute gradient using a NanoLC-Ultra 1D plus (Eksigent, Dublin CA) nano-pump and subsequently analyzed using a TripleTOF 5600 mass spectrometer (AB SCIEX, Concord, Ontario, Canada). The gradient was delivered at 200 nL/min, initiating from 2% acetonitrile with 0.1% formic acid to 35% acetonitrile with 0.1% formic acid over 90 minutes, followed by a 15-minute wash using 80% acetonitrile with 0.1% formic acid, and a 15-minute equilibration period to 2% acetonitrile with 0.1% formic acid. Instrument performance was monitored daily using a system suitability test data consisting of a 30-minute gradient injection of 30fm BSA and 60fm of casein protein digest (both standards were trypsin digested in-house from commercial protein stocks (Sigma)) that was run before each sample. Performance monitoring consisted of tracking Peak intensities, mass accuracies, and retention times to ensure LCMS data quality was consistent throughout the project. The mass accuracy of the 5600 instrument was calibrated before each sample analysis using an automated routine.

The instrument method used in DDA mode consisted of one 250 ms MS1 TOF survey scan from 400-1250 Da followed by isolation of the top 20 MS2 candidate ions for 100 ms per ion. Only ions charged 2+ to 4+ exceeding a threshold of 250 cps were considered for MS2, and former precursors were excluded for 15 s following isolation.

### Mass spectrometry data analysis

AB SCIEX WIFF MS files were first converted to mzXML using Proteowizard^90^ implemented in ProHits v4.0^91^. mzML and mzXML files were searched using Mascot (version 2.3.02) and Comet version 2012.02rev.0^92^ against the NCBI RefSeq database (version 57, January 30, 2013) containing a total of 72,482 human and adenovirus sequences supplemented with common contaminants from the Max Planck Institute (http://lotus1.gwdg.de/mpg/mmbc/maxquant_input.nsf/7994124a4298328fc125748d0048fee2/$ FILE/contaminants.fasta) and the Global Proteome Machine (GPM; https://www.thegpm.org/crap/index.html). The database parameters were set to search for tryptic cleavages, permitting up to two missed cleavage sites per peptide with a mass tolerance of 35 ppm for precursors with charges +2 to +4 and a tolerance of ± 0.15 amu for fragment ions. Asparagine or glutamine deamination and methionine oxidation were allowed as variable modifications. The results from each search engine were subsequently analyzed through the Trans-Proteomic Pipeline (version 4.6 Occupy rev 3) using the iProphet pipeline^93^.

### Significance analysis of INTeractome (SAINT) analysis

SAINTexpress (version 3.6.1)^94^ was used as a statistical tool to filter out likely contaminants. Briefly: all protein entries identified with ≥ 2 unique peptides and iProphet score ≥0.95 were used for SAINTexpress analysis, and biological duplicates were used for each bait. Negative controls [fusions of BirA∗-FLAG to EGFP or Nanoluciferase or untransfected HEK293 cells; 10 controls in total – Table S1, sheet B: Controls] were compressed to 5 virtual controls to maximize stringency in scoring, as previously described^95^. SAINTexpress analysis was performed using the default parameters, and only those entries passing a calculated Bayesian FDR (BFDR) ≤ 1% were considered high-confidence.

### Luminescence assays for SMAD2 characterization

For 3TP-lux reporter assays, MDA231 cells were seeded into clear 96-well plates at 15,000 cells per well in RPMI 1640 medium (Gibco) supplemented with 5% FBS growth medium. The following day, each well was transfected with 100 ng of 3TP-lux (Attisano Lab), 50 ng 3xFLAG-tagged ORF, and 5 ng Nanoluciferase with X-tremeGENE 9™ (MilliporeSigma) following the manufacturer’s protocol. The next day, the media was removed, and cells were starved with RPMI supplemented with 0.2% FBS for 8 hours. Then, the media was removed and replaced with 100 µl of 0.2% serum media +/-100 pM TGFβ (a kind gift from the Attisano lab, University of Toronto). After an overnight incubation (16 hours), cells were washed with 1 x PBS, lysed with 80 µl/well HENG buffer (20 mM HEPES-KOH pH 7.9, 150 mM NaCl, 2 mM EDTA pH 8.0, 20 mM Na2MoO4, 0.5% Triton X-100, 5% glycerol) containing protease inhibitors (1 μg/mL aprotinin, 1 μg/mL leupeptin, 1 μg/mL pepstatin, 0.5 mM PMSF) and incubated on a shaker at 4°C. After 20 minutes of incubation, 20 µl of the sample and 20 µl of Firefly Luciferase Lysis Buffer (150 mM Tris-HCL pH 8.0, 75 mM NaCl, 3 mM MgCl2, 0.25% Triton X-100, 15 mM DTT, 0.6 mM Coenzyme A, 0.45 mM ATP pH 7.0, 250 µg/mL D-luciferin) was added to a 96-well white opaque plate. Firefly signal was measured using a BioTek Synergy Neo microplate reader after the plate was incubated at room temperature for 10 minutes. Then, 20 µl of Firefly Stop & Glo Buffer (20 mM Tris-HCL pH 7.5, 150 mM KCl, 45 mM EDTA pH 8.0, 0.5% Tergitol NP9, 60 µM PTC124, 50 mM Thioacetamide, 5 µM Fuzimarine) was added to the 96-well plate, incubated for 10 minutes at room temperature, and Nanoluciferase signal was read.

For the LUMIER assay, 293T cells were seeded into clear 96-well plates at 30,000 cells per well in DMEM supplemented with 10% FBS. The following day, each well was transfected with 75 ng of 3xFLAG-tagged ORF (prey) and 75 ng of either Nanoluc-SMAD2 WT or Nanoluc-SMAD2-D304G in a mixture with 0.6 µL polyethylenimine (PEI) and 50 µL OptiMEM. After two days, cells were treated with 5 pM TGFβ for 5 minutes then cells were washed with 1xPBS, and then lysed with ice-cold HENG buffer (20 mM HEPES pH 7.9, 150 mM NaCl, 2mM EDTA pH 8, 0.5% Triton X-100, 5% glycerol) containing protease inhibitors (1 µg/ml aprotinin, 1 µg/ml leupeptin, 1µg/ml pepstatin, 0.2 mM PMSF). The lysates were then transferred from the 96-well plates into opaque white 384-well plates that were pre-coated with monoclonal anti-FLAG M2 antibody (Millipore Sigma, F1804). Plates were blocked with 1% BSA/5% sucrose/0.5% Tween 20/1xPBS. The 384-well plates were incubated for 3 hours at 4°C with mild shaking and then washed with HENG buffer (without protease inhibitors) using an automated plate washer. Luminescence was measured with a BioTek Synergy Neo microplate reader five minutes after adding furimazine luciferase reagent dissolved 1:200 in luciferase buffer (20 mM Tris-HCl pH 7.5, 1 mM EDTA, 150 mM KCl, 0.5% Tergitol NP9). Afterwards, HRP-conjugated anti-FLAG antibody (1:10,000, Millipore Sigma, A8592) diluted in ELISA buffer (1xPBS, 1% goat serum, 1% Tween 20) was added to wells and plates were incubated for 90 minutes at room temperature with mild shaking. Plates were then washed with 0.1% Tween 20/1xPBS using an automated plate washer. ELISA signal was measured using a BioTek Synergy Neo microplate reader one minute after adding SuperSignal ELISA Pico Chemiluminescent substrate (ThermoFisher Scientific, 37069).

### Quantification and statistical analysis

Statistical analysis was performed using GraphPad Prism. Statistical tests used and sample sizes are described in figure legends.

## SUPPLEMENTARY FIGURES

**Figure S1.**
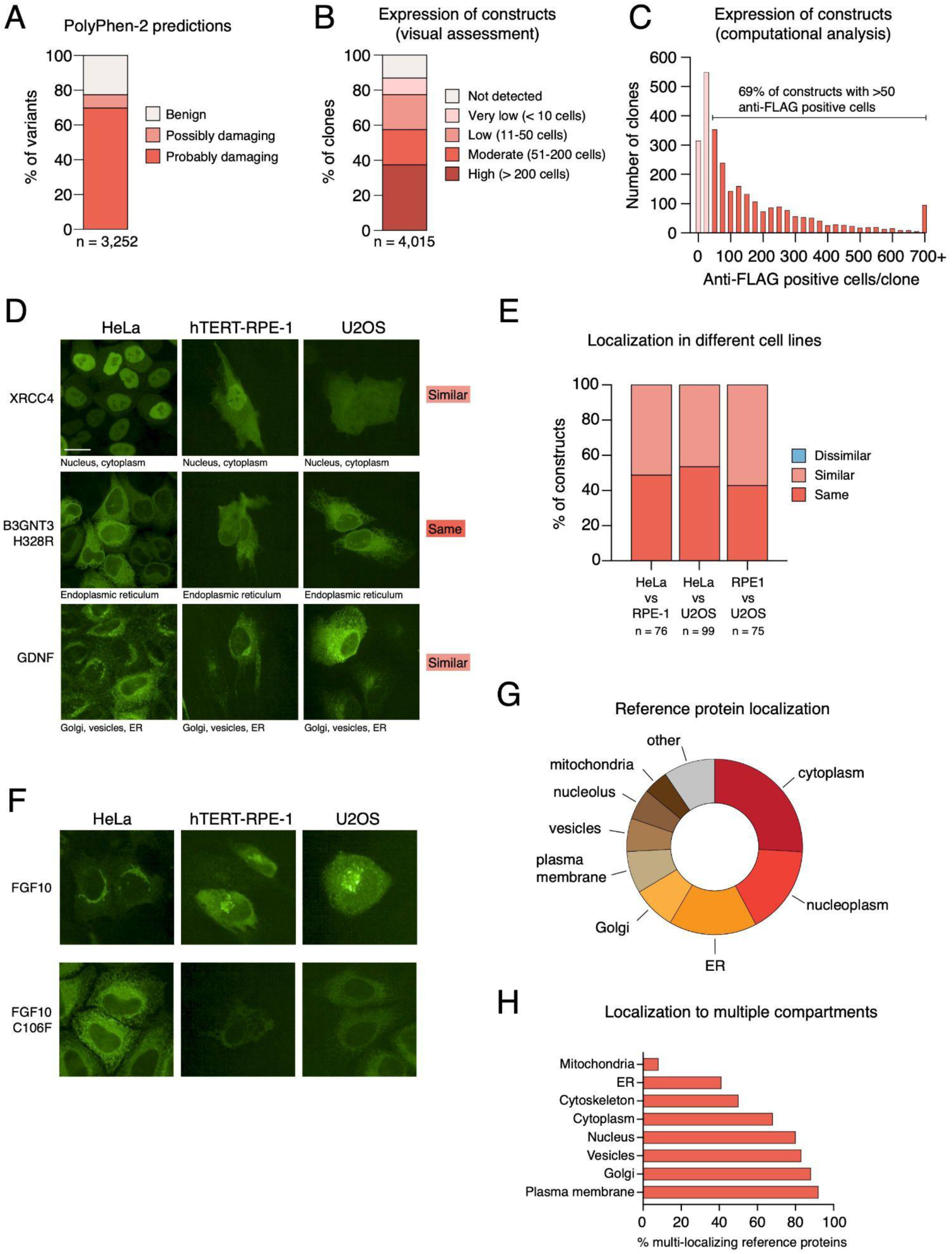
Characterization and quality control of the missense variant collection. Related to Figure **A)** Available PolyPhen-2 predictions for the variants used in this study. **B)** Expression of 3xFLAG-V5 tagged constructs by immunofluorescence as assessed by visual inspection. **C)** Expression of 3xFLAG-V5 tagged constructs by immunofluorescence as assessed by CellProfiler. **D)** Examples of localization patterns in HeLa, hTERT RPE-1, and U2OS cells. **E)** Comparison of localization patterns in HeLa, hTERT RPE-1, and U2OS cells. **F)** An example of similar variant mislocalization in HeLa, hTERT RPE-1, and U2OS cells. **G)** Localization patterns of transfected reference proteins. If protein was localized to multiple compartments, all compartments were included in the graph. **H)** Percentage of constructs localizing to multiple compartments.

**Figure S2.**
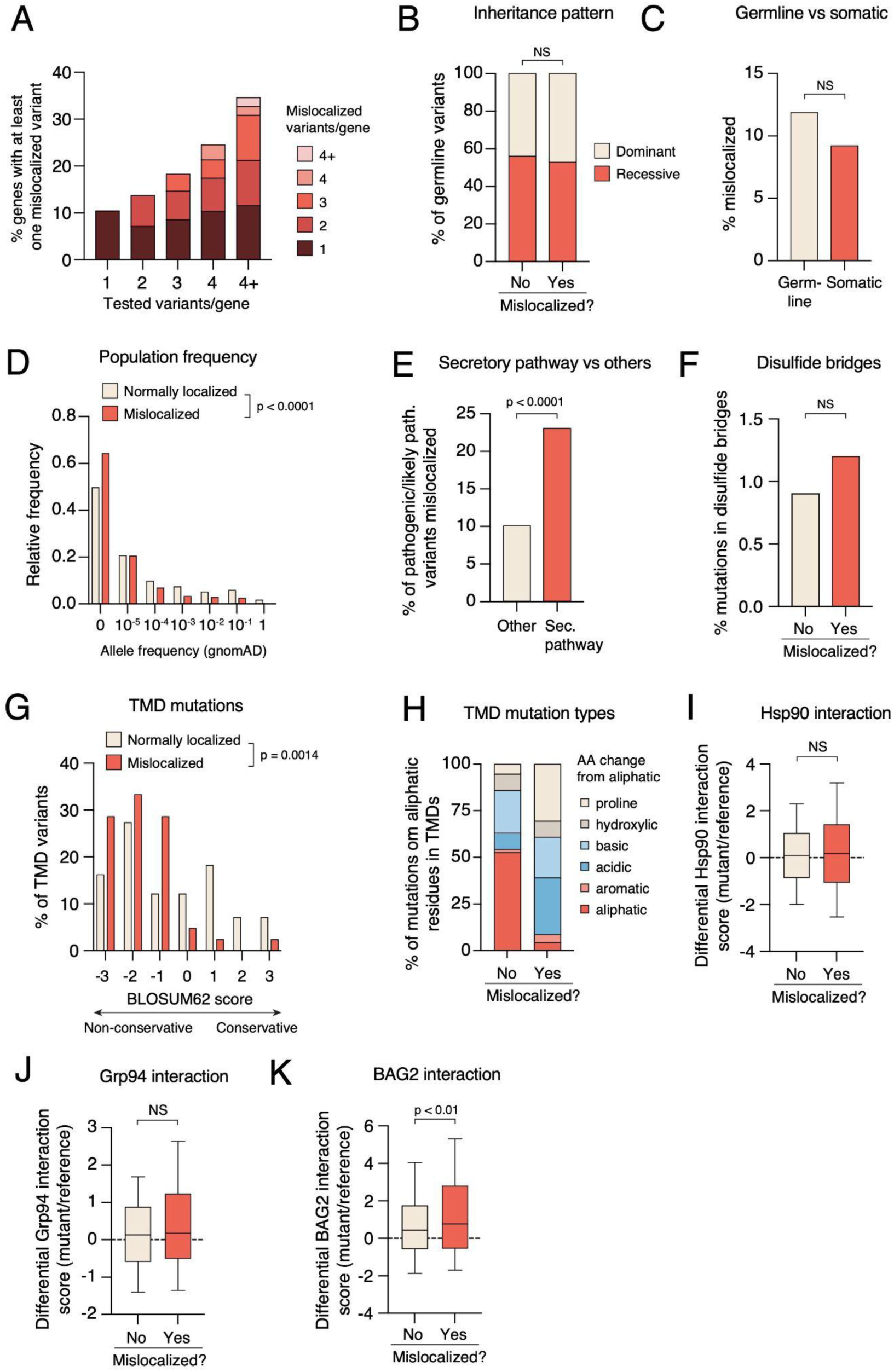
Features of mislocalized variants and proteins. Related to Figures 3 **and 4. A)** Fraction of genes with at least one mislocalized variant as a function of the number of variants tested. **B)** Annotated inheritance pattern of mislocalized variants. **C)** Mislocalization frequency for germline and somatic variants. **D)** Population frequencies of mislocalized and normally localized variants (extracted from gnomAD). **E)** Fraction of pathogenic/likely pathogenic variants that are mislocalized for variants of proteins trafficked through the secretory pathway and for other proteins. **F)** Frequency of mutations in cysteines forming disulfide bridges for normally localized and mislocalized variants. **G)** BLOSUM62 score for missense mutations in transmembrane domains for normally localized and mislocalized variants. **H)** Distribution of mutations of aliphatic amino acids (alanine, methionine, leucine, isoleucine, valine) to other amino acids in transmembrane domains for normally localized and mislocalized variants. **I-K)** Comparison of chaperone interactions of reference proteins and mislocalized or normally localized proteins for Hsp90/HSP90AB1 **(I)** and Grp94/HSP90B1 **(J)**, and BAG2 **(K).** Statistical significance was calculated with the Mann-Whitney test (panels D, G, I, J, K), and Fisher’s exact test (panel E).

**Figure S3.**
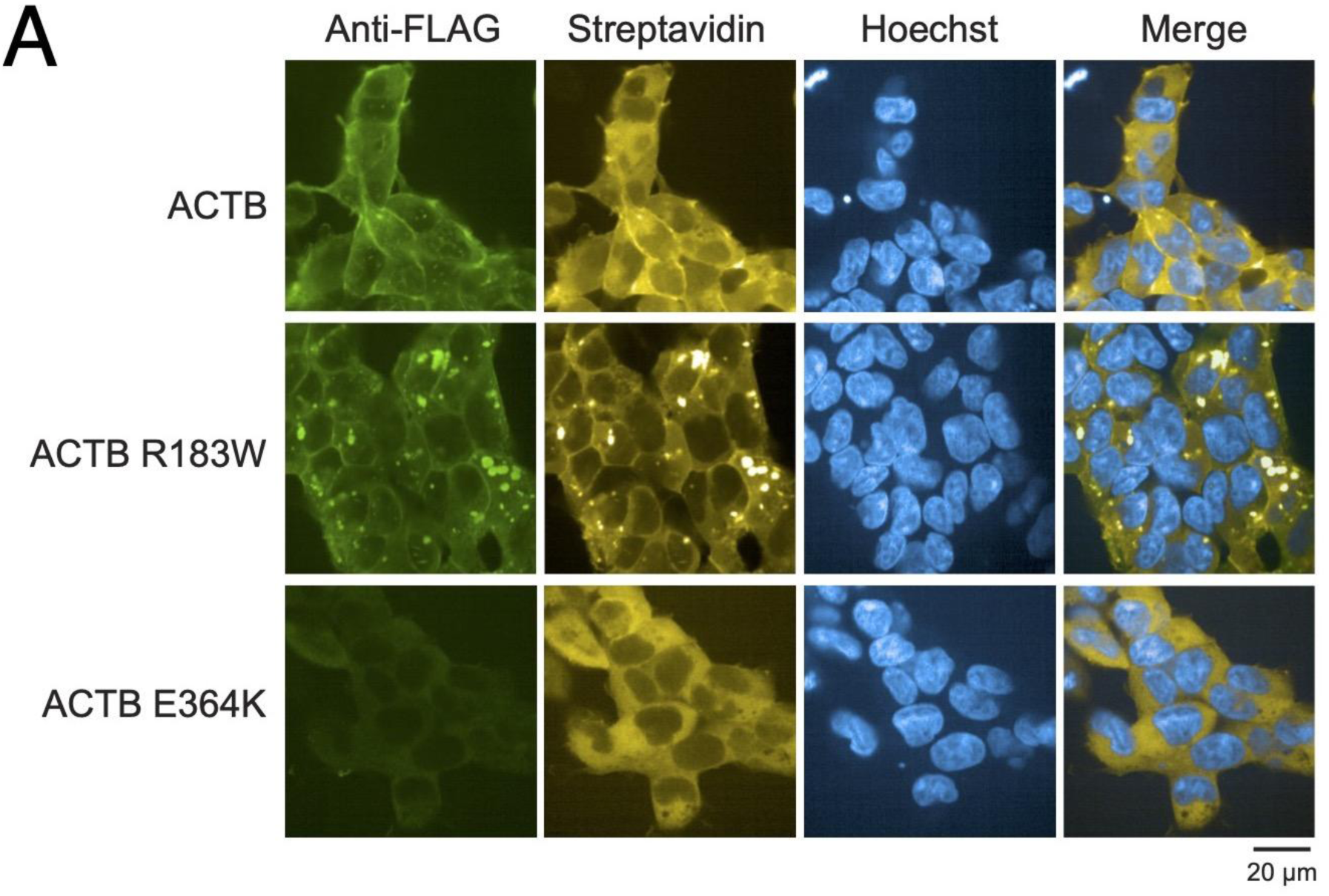
Localization of wild-type and mutant beta-actin constructs tagged with BirA2-FLAG in HEK293 cells. Stable HEK293 Flp-In T-REx cells were induced with tetracycline for 24 hours and treated with biotin for 12 hours prior to fixation and anti-FLAG (green), streptavidin (yellow), and Hoechst (blue) staining.

## REFERENCES 1

1. Tabet, D., Parikh, V., Mali, P., Roth, F.P., and Claussnitzer, M. (2022). Scalable Functional Assays for the Interpretation of Human Genetic Variation. Annu. Rev. Genet. 56, 441–465. 10.1146/annurev-genet-072920-032107.

2. Taipale, M. (2018). Disruption of protein function by pathogenic mutations: common and uncommon mechanisms. Biochem. Cell Biol. 37, 508.

3. Capriotti, E., Ozturk, K., and Carter, H. (2019). Integrating molecular networks with genetic variant interpretation for precision medicine. WIREs Syst. Biol. Med. 11. 10.1002/wsbm.1443.

4. Vihinen, M. (2021). Functional effects of protein variants. Biochimie 180, 104–120. 10.1016/j.biochi.2020.10.009.

5. Taverna, D.M., and Goldstein, R.A. (2001). Why are proteins marginally stable? Proteins 46, 105–109.

6. Matreyek, K.A., Starita, L.M., Stephany, J.J., Martin, B., Chiasson, M.A., Gray, V.E., Kircher, M., Khechaduri, A., Dines, J.N., Hause, R.J., et al. (2018). Multiplex assessment of protein variant abundance by massively parallel sequencing. Nat. Genet. 22, 925.

7. Sahni, N., Yi, S., Taipale, M., Fuxman Bass, J.I., Coulombe-Huntington, J., Yang, F., Peng, J., Weile, J., Karras, G.I., Wang, Y., et al. (2015). Widespread Macromolecular Interaction Perturbations in Human Genetic Disorders. Cell 161, 647–660.

8. Mosca, R., Tenorio-Laranga, J., Olivella, R., Alcalde, V., Ceol, A., Soler-Lόpez, M., and Aloy, P. (2015). dSysMap: exploring the edgetic role of disease mutations. Nat. Methods 12, 167–168.

9. Dang, L., White, D.W., Gross, S., Bennett, B.D., Bittinger, M.A., Driggers, E.M., Fantin, V.R., Jang, H.G., Jin, S., Keenan, M.C., et al. (2009). Cancer-associated IDH1 mutations produce 2-hydroxyglutarate. Nature 462, 739–744.

10. Barrera, L.A., Vedenko, A., Kurland, J.V., Rogers, J.M., Gisselbrecht, S.S., Rossin, E.J., Woodard, J., Mariani, L., Kock, K.H., Inukai, S., et al. (2016). Survey of variation in human transcription factors reveals prevalent DNA binding changes. Sci. N. Y. NY 351, 1450– 1454.

11. Sahni, N., Yi, S., Zhong, Q., Jailkhani, N., Charloteaux, B., Cusick, M.E., and Vidal, M. (2013). Edgotype: a fundamental link between genotype and phenotype. Curr. Opin. Genet. Span Classnocasespan Dev. 23, 649–657.

12. Zhong, Q., Simonis, N., Li, Q.-R., Charloteaux, B., Heuze, F., Klitgord, N., Tam, S., Yu, H., Venkatesan, K., Mou, D., et al. (2009). Edgetic perturbation models of human inherited disorders. Mol. Syst. Biol. 5, 321.

13. Wang, X., Wei, X., Thijssen, B., Das, J., Lipkin, S.M., and Yu, H. (2012). Three-dimensional reconstruction of protein networks provides insight into human genetic disease. Nat. Biotechnol. 30, 159–164.

14. Xiong, D., Zhao, J., Qiu, Y., Zhou, Y., Lee, D., Gupta, S., Lu, W., Liang, S., Kang, J.J., Eng, C., et al. (2023). 3D structural human interactome reveals proteome-wide perturbations by disease mutations. BioRxiv Prepr. Serv. Biol., 2023.04.24.538110. 10.1101/2023.04.24.538110.

15. Park, S., Yang, J.-S., Shin, Y.-E., Park, J., Jang, S.K., and Kim, S. (2011). Protein localization as a principal feature of the etiology and comorbidity of genetic diseases. Mol. Syst. Biol. 7, 494. 10.1038/msb.2011.29.

16. Hung, M.-C., and Link, W. (2011). Protein localization in disease and therapy. J. Cell Sci. 124, 3381–3392.

17. Laurila, K., and Vihinen, M. (2009). Prediction of disease-related mutations affecting protein localization. BMC Genomics 10, 122.

18. Mensah, M.A., Niskanen, H., Magalhaes, A.P., Basu, S., Kircher, M., Sczakiel, H.L., Reiter, A.M.V., Elsner, J., Meinecke, P., Biskup, S., et al. (2023). Aberrant phase separation and nucleolar dysfunction in rare genetic diseases. Nature. 10.1038/s41586-022-05682-1.

19. Cutting, G.R. (2015). Cystic fibrosis genetics: from molecular understanding to clinical application. Nat. Rev. Genet. 16, 45–56. 10.1038/nrg3849.

20. Lopes-Pacheco, M. (2019). CFTR Modulators: The Changing Face of Cystic Fibrosis in the Era of Precision Medicine. Front. Pharmacol. 10, 1662. 10.3389/fphar.2019.01662.

21. Hasle, N., Matreyek, K.A., and Fowler, D.M. (2019). The Impact of Genetic Variants on PTEN Molecular Functions and Cellular Phenotypes. Cold Spring Harb. Perspect. Med. 9, a036228. 10.1101/cshperspect.a036228.

22. Carpten, J.D., Faber, A.L., Horn, C., Donoho, G.P., Briggs, S.L., Robbins, C.M., Hostetter, G., Boguslawski, S., Moses, T.Y., Savage, S., et al. (2007). A transforming mutation in the pleckstrin homology domain of AKT1 in cancer. Nature 448, 439–444. 10.1038/nature05933.

23. Rodriguez, J.A., Au, W.W.Y., and Henderson, B.R. (2004). Cytoplasmic mislocalization of BRCA1 caused by cancer-associated mutations in the BRCT domain. Exp. Cell Res. 293, 14–21. 10.1016/j.yexcr.2003.09.027.

24. Falini, B., Nicoletti, I., Martelli, M.F., and Mecucci, C. (2007). Acute myeloid leukemia carrying cytoplasmic/mutated nucleophosmin (NPMc+ AML): biologic and clinical features. Blood 109, 874–885. 10.1182/blood-2006-07-012252.

25. Patel, A., Lee, H.O., Jawerth, L., Maharana, S., Jahnel, M., Hein, M.Y., Stoynov, S., Mahamid, J., Saha, S., Franzmann, T.M., et al. (2015). A Liquid-to-Solid Phase Transition of the ALS Protein FUS Accelerated by Disease Mutation. Cell 162, 1066–1077.

26. Stenson, P.D., Mort, M., Ball, E.V., Chapman, M., Evans, K., Azevedo, L., Hayden, M., Heywood, S., Millar, D.S., Phillips, A.D., et al. (2020). The Human Gene Mutation Database (HGMD®): optimizing its use in a clinical diagnostic or research setting. Hum. Genet. 139, 1197–1207. 10.1007/s00439-020-02199-3.

27. Taipale, M., Krykbaeva, I., Whitesell, L., Santagata, S., Zhang, J., Liu, Q., Gray, N.S., and Lindquist, S. (2013). Chaperones as thermodynamic sensors of drug-target interactions reveal kinase inhibitor specificities in living cells. Nat. Biotechnol. 31, 630–637.

28. Taipale, M., Tucker, G., Peng, J., Krykbaeva, I., Lin, Z.-Y., Larsen, B., Choi, H., Berger, B., Gingras, A.-C., and Lindquist, S. (2014). A quantitative chaperone interaction network reveals the architecture of cellular protein homeostasis pathways. Cell 158, 434–448.

29. Kim, E., Ilic, N., Shrestha, Y., Zou, L., Kamburov, A., Zhu, C., Yang, X., Lubonja, R., Tran, N., Nguyen, C., et al. (2016). Systematic Functional Interrogation of Rare Cancer Variants Identifies Oncogenic Alleles. Cancer Discov. 6, 714–726.

30. Adzhubei, I.A., Schmidt, S., Peshkin, L., Ramensky, V.E., Gerasimova, A., Bork, P., Kondrashov, A.S., and Sunyaev, S.R. (2010). A method and server for predicting damaging missense mutations. Nat. Methods 7, 248–249.

31. Landrum, M.J., Lee, J.M., Benson, M., Brown, G., Chao, C., Chitipiralla, S., Gu, B., Hart, J., Hoffman, D., Hoover, J., et al. (2016). ClinVar: public archive of interpretations of clinically relevant variants. Nucleic Acids Res. 44, D862–8.

32. Stirling, D.R., Swain-Bowden, M.J., Lucas, A.M., Carpenter, A.E., Cimini, B.A., and Goodman, A. (2021). CellProfiler 4: improvements in speed, utility and usability. BMC Bioinformatics 22, 433. 10.1186/s12859-021-04344-9.

33. Thul, P.J., Åkesson, L., Wiking, M., Mahdessian, D., Geladaki, A., Ait Blal, H., Alm, T., Asplund, A., Björk, L., Breckels, L.M., et al. (2017). A subcellular map of the human proteome. Sci. N. Y. NY 356, eaal3321.

34. Christoforou, A., Mulvey, C.M., Breckels, L.M., Geladaki, A., Hurrell, T., Hayward, P.C., Naake, T., Gatto, L., Viner, R., Arias, A.M., et al. (2016). A draft map of the mouse pluripotent stem cell spatial proteome. Nat. Commun. 7, 9992. 10.1038/ncomms9992.

35. Itzhak, D.N., Tyanova, S., Cox, J., and Borner, G.H. (2016). Global, quantitative and dynamic mapping of protein subcellular localization. eLife 5, e16950. 10.7554/eLife.16950.

36. Orre, L.M., Vesterlund, M., Pan, Y., Arslan, T., Zhu, Y., Fernandez Woodbridge, A., Frings, O., Fredlund, E., and Lehtiö, J. (2019). SubCellBarCode: Proteome-wide Mapping of Protein Localization and Relocalization. Mol. Cell 73, 166–182.e7. 10.1016/j.molcel.2018.11.035.

37. Go, C.D., Knight, J.D.R., Rajasekharan, A., Rathod, B., Hesketh, G.G., Abe, K.T., Youn, J.-Y., Samavarchi-Tehrani, P., Zhang, H., Zhu, L.Y., et al. (2021). A proximity-dependent biotinylation map of a human cell. Nature 595, 120–124. 10.1038/s41586-021-03592-2.

38. Stadler, C., Rexhepaj, E., Singan, V.R., Murphy, R.F., Pepperkok, R., Uhlen, M., Simpson, J.C., and Lundberg, E. (2013). Immunofluorescence and fluorescent-protein tagging show high correlation for protein localization in mammalian cells. Nat. Methods 10, 315–323.

39. Chong, Y.T., Koh, J.L.Y., Friesen, H., Duffy, K., Cox, M.J., Moses, A., Moffat, J., Boone, C., and Andrews, B.J. (2015). Yeast Proteome Dynamics from Single Cell Imaging and Automated Analysis. Cell 161, 1413–1424.

40. Sun, Z., and Brodsky, J.L. (2019). Protein quality control in the secretory pathway. J. Cell Biol. 218, 3171–3187. 10.1083/jcb.201906047.

41. Wang, X., Osinska, H., Klevitsky, R., Gerdes, A.M., Nieman, M., Lorenz, J., Hewett, T., and Robbins, J. (2001). Expression of R120G-alphaB-crystallin causes aberrant desmin and alphaB-crystallin aggregation and cardiomyopathy in mice. Circ. Res. 89, 84–91. 10.1161/hh1301.092688.

42. Qi, L.-B., Hu, L.-D., Liu, H., Li, H.-Y., Leng, X.-Y., and Yan, Y.-B. (2016). Cataract-causing mutation S228P promotes βB1-crystallin aggregation and degradation by separating two interacting loops in C-terminal domain. Protein Cell 7, 501–515. 10.1007/s13238-016-0284-3.

43. Marzahn, M.R., Marada, S., Lee, J., Nourse, A., Kenrick, S., Zhao, H., Ben-Nissan, G., Kolaitis, R.-M., Peters, J.L., Pounds, S., et al. (2016). Higher-order oligomerization promotes localization of SPOP to liquid nuclear speckles. EMBO J. 35, 1254–1275. 10.15252/embj.201593169.

44. Usher, E.T., Sabri, N., Rohac, R., Boal, A.K., Mittag, T., and Showalter, S.A. (2021). Intrinsically disordered substrates dictate SPOP subnuclear localization and ubiquitination activity. J. Biol. Chem. 296, 100693. 10.1016/j.jbc.2021.100693.

45. Janouskova, H., El Tekle, G., Bellini, E., Udeshi, N.D., Rinaldi, A., Ulbricht, A., Bernasocchi, T., Civenni, G., Losa, M., Svinkina, T., et al. (2017). Opposing effects of cancer-type-specific SPOP mutants on BET protein degradation and sensitivity to BET inhibitors. Nat. Med. 23, 1046–1054. 10.1038/nm.4372.

46. Banani, S.F., Afeyan, L.K., Hawken, S.W., Henninger, J.E., Dall’Agnese, A., Clark, V.E., Platt, J.M., Oksuz, O., Hannett, N.M., Sagi, I., et al. (2022). Genetic variation associated with condensate dysregulation in disease. Dev. Cell 57, 1776–1788.e8. 10.1016/j.devcel.2022.06.010.

47. Karczewski, K.J., Francioli, L.C., Tiao, G., Cummings, B.B., Alföldi, J., Wang, Q., Collins, R.L., Laricchia, K.M., Ganna, A., Birnbaum, D.P., et al. (2020). The mutational constraint spectrum quantified from variation in 141,456 humans. Nature 581, 434–443. 10.1038/s41586-020-2308-7.

48. de Nijs, L., Wolkoff, N., Coumans, B., Delgado-Escueta, A.V., Grisar, T., and Lakaye, B. (2012). Mutations of EFHC1, linked to juvenile myoclonic epilepsy, disrupt radial and tangential migrations during brain development. Hum. Mol. Genet. 21, 5106–5117. 10.1093/hmg/dds356.

49. Bailey, J.N., Patterson, C., de Nijs, L., Durón, R.M., Nguyen, V.-H., Tanaka, M., Medina, M.T., Jara-Prado, A., Martínez-Juárez, I.E., Ochoa, A., et al. (2017). EFHC1 variants in juvenile myoclonic epilepsy: reanalysis according to NHGRI and ACMG guidelines for assigning disease causality. Genet. Med. Off. J. Am. Coll. Med. Genet. 19, 144–156. 10.1038/gim.2016.86.

50. Hosagrahara, V.P., Rettie, A.E., Hassett, C., and Omiecinski, C.J. (2004). Functional analysis of human microsomal epoxide hydrolase genetic variants. Chem. Biol. Interact. 150, 149–159. 10.1016/j.cbi.2004.07.004.

51. Sile, S., Gillani, N.B., Velez, D.R., Vanoye, C.G., Yu, C., Byrne, L.M., Gainer, J.V., Brown, N.J., Williams, S.M., and George, A.L. (2007). Functional BSND variants in essential hypertension. Am. J. Hypertens. 20, 1176–1182. 10.1016/j.amjhyper.2007.07.003.

52. Koga, M., Ishiguro, H., Yazaki, S., Horiuchi, Y., Arai, M., Niizato, K., Iritani, S., Itokawa, M., Inada, T., Iwata, N., et al. (2009). Involvement of SMARCA2/BRM in the SWI/SNF chromatin-remodeling complex in schizophrenia. Hum. Mol. Genet. 18, 2483–2494. 10.1093/hmg/ddp166.

53. Katano, M., Numata, T., Aguan, K., Hara, Y., Kiyonaka, S., Yamamoto, S., Miki, T., Sawamura, S., Suzuki, T., Yamakawa, K., et al. (2012). The juvenile myoclonic epilepsy-related protein EFHC1 interacts with the redox-sensitive TRPM2 channel linked to cell death. Cell Calcium 51, 179–185. 10.1016/j.ceca.2011.12.011.

54. Fragoza, R., Das, J., Wierbowski, S.D., Liang, J., Tran, T.N., Liang, S., Beltran, J.F. Rivera-Erick, C.A., Ye, K., Wang, T.-Y., et al. (2019). Extensive disruption of protein interactions by genetic variants across the allele frequency spectrum in human populations. Nat. Commun. 10, 4141. 10.1038/s41467-019-11959-3.

55. Toriyama, M., Mizuno, N., Fukami, T., Iguchi, T., Toriyama, M., Tago, K., and Itoh, H. (2012). Phosphorylation of doublecortin by protein kinase A orchestrates microtubule and actin dynamics to promote neuronal progenitor cell migration. J. Biol. Chem. 287, 12691– 12702. 10.1074/jbc.M111.316307.

56. Ori-McKenney, K.M., McKenney, R.J., Huang, H.H., Li, T., Meltzer, S., Jan, L.Y., Vale, R.D., Wiita, A.P., and Jan, Y.N. (2016). Phosphorylation of β-Tubulin by the Down Syndrome Kinase, Minibrain/DYRK1a, Regulates Microtubule Dynamics and Dendrite Morphogenesis. Neuron 90, 551–563. 10.1016/j.neuron.2016.03.027.

57. Kunze, M., and Berger, J. (2015). The similarity between N-terminal targeting signals for protein import into different organelles and its evolutionary relevance. Front. Physiol. 6, 259. 10.3389/fphys.2015.00259.

58. Zhang, B., Spreafico, M., Zheng, C., Yang, A., Platzer, P., Callaghan, M.U., Avci, Z., Ozbek, N., Mahlangu, J., Haw, T., et al. (2008). Genotype-phenotype correlation in combined deficiency of factor V and factor VIII. Blood 111, 5592–5600. 10.1182/blood-2007-10-113951.

59. Taipale, M., Jarosz, D.F., and Lindquist, S. (2010). HSP90 at the hub of protein homeostasis: emerging mechanistic insights. Nat. Rev. Mol. Cell Biol. 11, 515–528.

60. Rosenzweig, R., Nillegoda, N.B., Mayer, M.P., and Bukau, B. (2019). The Hsp70 chaperone network. Nat. Rev. Mol. Cell Biol. 20, 665–680. 10.1038/s41580-019-0133-3.

61. Wolf, N.I., van Spaendonk, R.M., Hobson, G.M., and Kamholz, J. (1993). PLP1 Disorders. In GeneReviews®, M. P. Adam, G. M. Mirzaa, R. A. Pagon, S. E. Wallace, L. J. Bean, K. W. Gripp, and A. Amemiya, eds. (University of Washington, Seattle).

62. Hagemann, T.L. (2022). Alexander disease: models, mechanisms, and medicine. Curr. Opin. Neurobiol. 72, 140–147. 10.1016/j.conb.2021.10.002.

63. Koizume, S., Takizawa, S., Fujita, K., Aida, N., Yamashita, S., Miyagi, Y., and Osaka, H. (2006). Aberrant trafficking of a proteolipid protein in a mild Pelizaeus-Merzbacher disease. Neuroscience 141, 1861–1869. 10.1016/j.neuroscience.2006.05.067.

64. Thomson, C.E., Montague, P., Jung, M., Nave, K.A., and Griffiths, I.R. (1997). Phenotypic severity of murine Plp mutants reflects in vivo and in vitro variations in transport of PLP isoproteins. Glia 20, 322–332. 10.1002/(sici)1098-1136(199708)20:4<322::aid-glia5>3.0.co;2-7.

65. Procaccio, V., Salazar, G., Ono, S., Styers, M.L., Gearing, M., Davila, A., Jimenez, R., Juncos, J., Gutekunst, C.-A., Meroni, G., et al. (2006). A mutation of beta -actin that alters depolymerization dynamics is associated with autosomal dominant developmental malformations, deafness, and dystonia. Am. J. Hum. Genet. 78, 947–960. 10.1086/504271.

66. Nunoi, H., Yamazaki, T., Tsuchiya, H., Kato, S., Malech, H.L., Matsuda, I., and Kanegasaki, S. (1999). A heterozygous mutation of beta-actin associated with neutrophil dysfunction and recurrent infection. Proc. Natl. Acad. Sci. U. S. A. 96, 8693–8698. 10.1073/pnas.96.15.8693.

67. Hundt, N., Preller, M., Swolski, O., Ang, A.M., Mannherz, H.G., Manstein, D.J., and Müller M. (2014). Molecular mechanisms of disease-related human β-actin mutations p.R183W and p.E364K. FEBS J. *281*, 5279–5291. 10.1111/febs.13068.

68. Rommelaere, H., Waterschoot, D., Neirynck, K., Vandekerckhove, J., and Ampe, C. (2004). A method for rapidly screening functionality of actin mutants and tagged actins. Biol. Proced. Online 6, 235–249. 10.1251/bpo94.

69. Gestaut, D., Roh, S.H., Ma, B., Pintilie, G., Joachimiak, L.A., Leitner, A., Walzthoeni, T., Aebersold, R., Chiu, W., and Frydman, J. (2019). The Chaperonin TRiC/CCT Associates with Prefoldin through a Conserved Electrostatic Interface Essential for Cellular Proteostasis. Cell 177, 751–765.e15. 10.1016/j.cell.2019.03.012.

70. Pollard, T.D. (2016). Actin and Actin-Binding Proteins. Cold Spring Harb. Perspect. Biol. 8, a018226. 10.1101/cshperspect.a018226.

71. Batlle, E., and Massagué, J. (2019). Transforming Growth Factor-β Signaling in Immunity and Cancer. Immunity 50, 924–940. 10.1016/j.immuni.2019.03.024.

72. Barrios-Rodiles, M., Brown, K.R., Ozdamar, B., Bose, R., Liu, Z., Donovan, R.S., Shinjo, F., Liu, Y., Dembowy, J., Taylor, I.W., et al. (2005). High-throughput mapping of a dynamic signaling network in mammalian cells. Sci. N. Y. NY *307*, 1621–1625.

73. Wrana, J.L., Attisano, L., Cárcamo, J., Zentella, A., Doody, J., Laiho, M., Wang, X.F., and Massagué, J. (1992). TGF beta signals through a heteromeric protein kinase receptor complex. Cell 71, 1003–1014. 10.1016/0092-8674(92)90395-s.

74. Uebel, C., and Phillips, C. (2019). Phase-separated protein dynamics are affected by fluorescent tag choice. MicroPublication Biol. 2019. 10.17912/micropub.biology.000143.

75. Kuang, D., Weile, J., Li, R., Ouellette, T.W., Barber, J.A., and Roth, F.P. (2020). MaveQuest: a web resource for planning experimental tests of human variant effects. Bioinforma. Oxf. Engl. *36*, 3938–3940. 10.1093/bioinformatics/btaa228.

76. Caicedo, J.C., Cooper, S., Heigwer, F., Warchal, S., Qiu, P., Molnar, C., Vasilevich, A.S., Barry, J.D., Bansal, H.S., Kraus, O., et al. (2017). Data-analysis strategies for image-based cell profiling. Nat. Methods 14, 849–863.

77. Rohban, M.H., Singh, S., Wu, X., Berthet, J.B., Bray, M.-A., Shrestha, Y., Varelas, X., Boehm, J.S., and Carpenter, A.E. (2017). Systematic morphological profiling of human gene and allele function via Cell Painting. eLife 6, e24060. 10.7554/eLife.24060.

78. Funk, L., Su, K.-C., Ly, J., Feldman, D., Singh, A., Moodie, B., Blainey, P.C., and Cheeseman, I.M. (2022). The phenotypic landscape of essential human genes. Cell 185, 4634–4653.e22. 10.1016/j.cell.2022.10.017.

79. Feldman, D., Singh, A., Schmid-Burgk, J.L., Carlson, R.J., Mezger, A., Garrity, A.J., Zhang, F., and Blainey, P.C. (2019). Optical Pooled Screens in Human Cells. Cell 179, 787–799.e17. 10.1016/j.cell.2019.09.016.

80. Van Goor, F., Straley, K.S., Cao, D., González, J., Hadida, S., Hazlewood, A., Joubran, J., Knapp, T., Makings, L.R., Miller, M., et al. (2006). Rescue of DeltaF508-CFTR trafficking and gating in human cystic fibrosis airway primary cultures by small molecules. Am. J. Physiol. Lung Cell. Mol. Physiol. 290, L1117–1130. 10.1152/ajplung.00169.2005.

81. Van Goor, F., Hadida, S., Grootenhuis, P.D.J., Burton, B., Stack, J.H., Straley, K.S., Decker, C.J., Miller, M., McCartney, J., Olson, E.R., et al. (2011). Correction of the F508del-CFTR protein processing defect in vitro by the investigational drug VX-809. Proc. Natl. Acad. Sci. U. S. A. 108, 18843–18848. 10.1073/pnas.1105787108.

82. Wang, M., Cotter, E., Wang, Y.-J., Fu, X., Whittsette, A.L., Lynch, J.W., Wiseman, R.L., Kelly, J.W., Keramidas, A., and Mu, T.-W. (2022). Pharmacological activation of ATF6 remodels the proteostasis network to rescue pathogenic GABAA receptors. Cell Biosci. 12, 48. 10.1186/s13578-022-00783-w.

83. Grandjean, J.M.D., and Wiseman, R.L. (2020). Small molecule strategies to harness the unfolded protein response: where do we go from here? J. Biol. Chem. 295, 15692–15711. 10.1074/jbc.REV120.010218.

84. Plate, L., Cooley, C.B., Chen, J.J., Paxman, R.J., Gallagher, C.M., Madoux, F., Genereux, J.C., Dobbs, W., Garza, D., Spicer, T.P., et al. (2016). Small molecule proteostasis regulators that reprogram the ER to reduce extracellular protein aggregation. eLife 5, e15550. 10.7554/eLife.15550.

85. Sun, S., Wang, C., Zhao, P., Kline, G.M., Grandjean, J.M.D., Jiang, X., Labaudiniere, R., Wiseman, R.L., Kelly, J.W., and Balch, W.E. (2023). Capturing the conversion of the pathogenic alpha-1-antitrypsin fold by ATF6 enhanced proteostasis. Cell Chem. Biol. 30, 22–42.e5. 10.1016/j.chembiol.2022.12.004.

86. Tunyasuvunakool, K., Adler, J., Wu, Z., Green, T., Zielinski, M., Žídek, A., Bridgland, A., Cowie, A., Meyer, C., Laydon, A., et al. (2021). Highly accurate protein structure prediction for the human proteome. Nature 596, 590–596. 10.1038/s41586-021-03828-1.

87. Hornbeck, P.V., Zhang, B., Murray, B., Kornhauser, J.M., Latham, V., and Skrzypek, E. (2015). PhosphoSitePlus, 2014: mutations, PTMs and recalibrations. Nucleic Acids Res. 43, D512-520. 10.1093/nar/gku1267.

88. Krassowski, M., Paczkowska, M., Cullion, K., Huang, T., Dzneladze, I., Ouellette, B.F.F., Yamada, J.T., Fradet-Turcotte, A., and Reimand, J. (2018). ActiveDriverDB: human disease mutations and genome variation in post-translational modification sites of proteins. Nucleic Acids Res. 46, D901–D910. 10.1093/nar/gkx973.

89. Couzens, A.L., Knight, J.D.R., Kean, M.J., Teo, G., Weiss, A., Dunham, W.H., Lin, Z.-Y., Bagshaw, R.D., Sicheri, F., Pawson, T., et al. (2013). Protein Interaction Network of the Mammalian Hippo Pathway Reveals Mechanisms of Kinase-Phosphatase Interactions. Sci. Signal. 6, rs15.

90. Adusumilli, R., and Mallick, P. (2017). Data Conversion with ProteoWizard msConvert. Methods Mol. Biol. Clifton NJ 1550, 339–368. 10.1007/978-1-4939-6747-6_23.

91. Liu, G., Knight, J.D.R., Zhang, J.P., Tsou, C.-C., Wang, J., Lambert, J.-P., Larsen, B., Tyers, M., Raught, B., Bandeira, N., et al. (2016). Data Independent Acquisition analysis in ProHits 4.0. J. Proteomics 149, 64–68. 10.1016/j.jprot.2016.04.042.

92. Eng, J.K., Jahan, T.A., and Hoopmann, M.R. (2013). Comet: an open-source MS/MS sequence database search tool. PROTEOMICS 13, 22–24.

93. Shteynberg, D., Deutsch, E.W., Lam, H., Eng, J.K., Sun, Z., Tasman, N., Mendoza, L., Moritz, R.L., Aebersold, R., and Nesvizhskii, A.I. (2011). iProphet: multi-level integrative analysis of shotgun proteomic data improves peptide and protein identification rates and error estimates. Mol. Span Classnocasespan Cell. Proteomics MCP 10, M111.007690.

94. Teo, G., Liu, G., Zhang, J., Nesvizhskii, A.I., Gingras, A.-C., and Choi, H. (2014). SAINTexpress: Improvements and additional features in Significance Analysis of INTeractome software. J. Proteomics 100, 37–43. 10.1016/j.jprot.2013.10.023.

95. Mellacheruvu, D., Wright, Z., Couzens, A.L., Lambert, J.-P., St-Denis, N.A., Li, T., Miteva, Y.V., Hauri, S., Sardiu, M.E., Low, T.Y., et al. (2013). The CRAPome: a contaminant repository for affinity purification-mass spectrometry data. Nat. Methods 10, 730–736.

